# Cerebral Organoids with Integrated Endothelial Networks Emulate the Neurovascular Unit and Mitigate Core Necrosis

**DOI:** 10.1101/2025.04.23.650161

**Authors:** Josep Fumadó Navarro, Siobhan Crilly, Wai Kit Chan, Shane Browne, John O. Mason, Catalina Vallejo Giraldo, Abhay Pandit, Mihai Lomora

## Abstract

Cerebral organoids (COs) are multicellular, self-organized, in vitro, 3D brain-like tissues used for developmental biology, disease modelling and drug screening. However, their lack of vascularity renders them less physiologically accurate. Vascularization of COs remains challenging due to the different requirements between COs and vascular cells, limited vascular network penetration within the organoid, and the absence of luminal perfusion. Here, we devised an encapsulation approach in which human brain microvascular endothelial cells (HBMVECs) were delivered to developing COs from progressively degrading extracellular matrix (ECM)-based hydrogel droplets. By tuning this hydrogel concentration and media composition, we observed enhanced vascular-like network formation that expanded within the organoid tissue. Using pathway inhibitors, we showed that a subset of the endothelial cells (ECs) originated from the CO itself, promoting network integration. Endothelial networks displayed blood-brain barrier (BBB) features, including astrocytic end-footlike interactions, pericyte wrapping, and collagen-laminin basal lamina. Vascularized COs exhibited greater media internalization and up to three-fold lower apoptosis than non-vascularized COs. This comprehensive 3D neurovascular model is a promising platform for cerebrovascular research and drug testing applications.

Graphical abstract

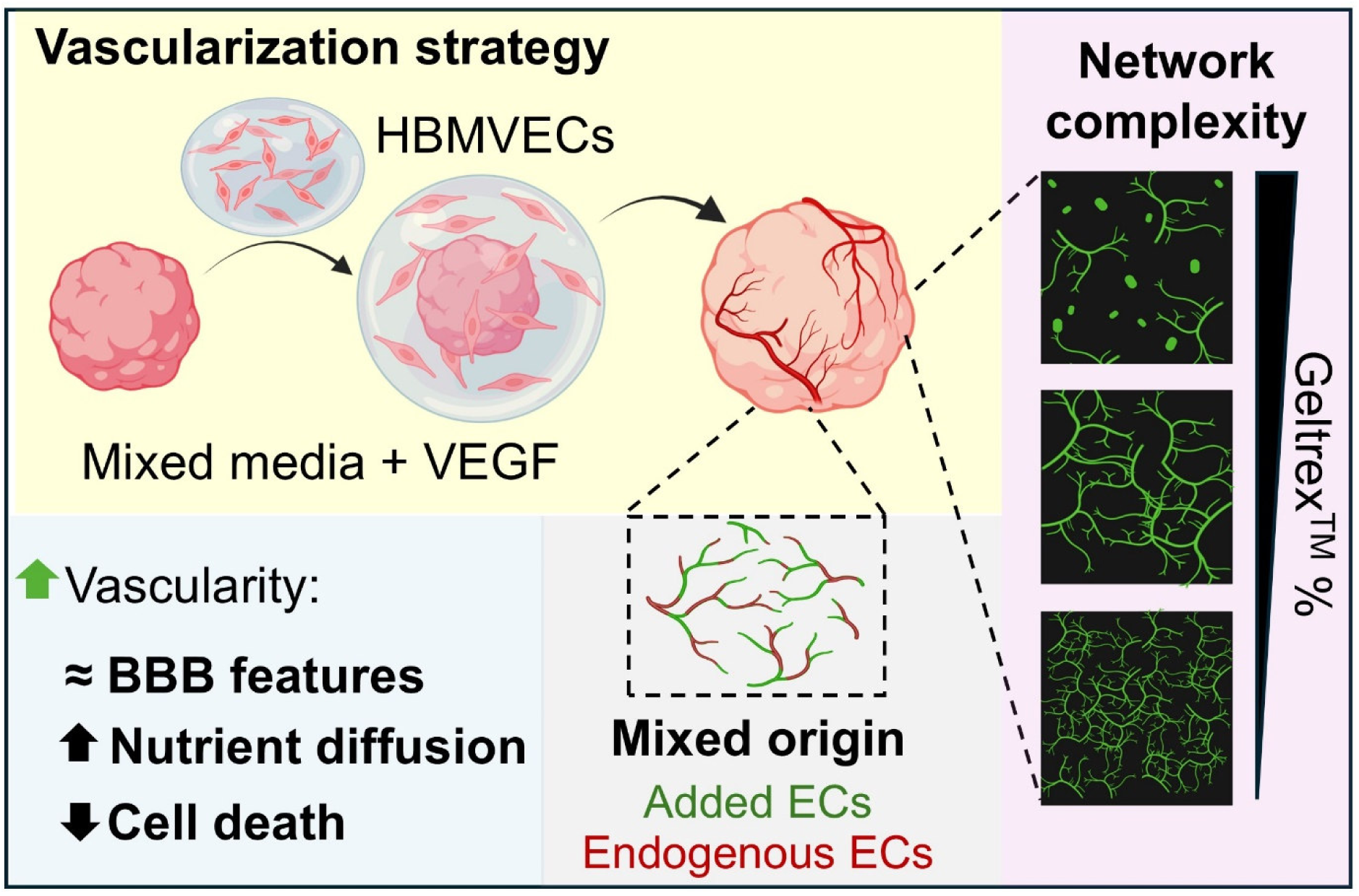

**Highlights:**

- **Addition and angiogenic stimulation of human brain microvascular endothelial cells (HBMVECs) co-cultured with cerebral organoids (COs) generate multicellular vascular-like networks**
- **Vascularization induces changes in organoid morphology but not on tissue stiffness**
- **Endothelial networks’ morphological features are correlated with the concentration of the supporting matrix**
- **A number of the endothelial cells (ECs) in the networks originate from the organoid itself**
- **Endothelial networks integrate within the organoid tissue interacting with astrocytes and pericyte-like cells, and are surrounded by basement membrane-like depositions**
- **Vascularized COs exhibit higher media diffusion, and a reduced necrotic core compared to non-vascularized COs**

## INTRODUCTION

Organoids have emerged as new high-throughput *in vitro* human models that have many applications in the biomedical field^1–4^. Cerebral (brain) organoids (COs) are self-organized tissue-like 3D cultures that resemble embryonic brain morphology and physiology^5,6^. They are obtained by aggregating induced pluripotent stem cells (iPSCs) or human embryonic stem cells (hESCs) followed by an orchestrated differentiation^7–10^. To date, several protocols for generating brain organoids are available to promote specific brain regions^11–16^. For instance, dorsal forebrain or cortical organoids are generated by inhibiting the bone morphogenic protein (BMP) and transforming growth factor beta (TGFβ) pathways (dual SMAD inhibition)^17,18^. These microphysical systems (MPS) have proven useful for neurodevelopmental studies, drug discovery, and understanding pathological mechanisms and have been extensively reviewed elsewhere^19–22^. However, one of the main limitations of cerebral organoids is their lack of vascularity^23^. As the organoid grows to millimetric sizes, the absence of vascularity leads to poor diffusion of nutrients and oxygen to the centre of the tissue causing hypoxia, metabolic stress and cell death, creating a necrotic core^24,25^. Additionally, vascularity promotes neural and glial differentiation and contributes to ventricular shaping and morphology of the developing brain^26^.

The cerebrovasculature has unique properties^27,28^, including a specific cell arrangement known as the neurovascular unit (NVU). This functional assembly consists of a tube of ECs wrapped by pericytes and astrocytic end-feet, and is structurally supported by a laminin– and collagen IV-rich basement membrane^29^. Together, this cellular coupling creates the BBB, a highly specialized and selective barrier that tightly regulates molecular transport to protect the brain from toxins, pathogens, and inflammatory cues circulating in the blood^30,31^.The low permeability of the BBB is obtained by a high expression of endothelial-specific adhesion molecules (i.e. platelet and endothelial cell adhesion molecule 1 or PECAM-1/CD31, and vascular endothelial cadherin or VE-CAD) and tight junctions (i.e., occludins and claudins)^32^. Given the critical role of the neurovascular unit in the anatomy and physiology of the developing and adult human brain, it is essential to recapitulate this highly organized cellular structure in a model to enhance its translational relevance for studying neurological and cerebrovascular disorders.

During embryonic development, neural and vascular tissues originate from different germinal layers (ectoderm and mesoderm, respectively) through different, yet highly coordinated, signaling pathways^33,34^. Thus, the challenge of vascularizing brain organoids lies in promoting vascular processes, such as vasculogenesis and angiogenesis, without disrupting neural tissue formation. Using genetic engineering (Ets variant 2 or ETV2 overexpression)^35^ and biochemical cues (i.e., vascular endothelial growth factor or VEGF, WNT)^36^, it has been possible to differentiate ECs from cerebral organoids and generate networks. Another strategy involves the direct addition of ECs (i.e., human umbilical vein endothelial cells/HUVECs or iPSC-derived ECs)^37–39^, together with mesodermal precursors and/or mural cells to cerebral organoids^40–43^, or fusing them with blood vessel organoids already containing all these vascular cell types^44,45^. The final *in vitro* approach involves employing advanced bioengineering techniques, such as bioprinting and microfluidic chips, to create perfusable vascular networks that serve as a platform where the organoids are allowed to interact with the vascular component^46–49^. Despite the achievements in these studies some limitations remain to be addressed. Models show variability, endothelial networks are superficial to the organoid tissue and achieving intra-organoid perfusion is extremely challenging, limiting vessel functionality.

Therefore, improving vascular integration in COs is crucial for reducing the necrotic core, promoting maturation, and better recapitulating *in vivo* tissue. In this study, we developed and characterized a strategy to generate vascular-like networks within COs, by adapting a commercially available protocol based on Lancaster et al^9^ (Figure 1). Modifications were introduced from day 8 onward, when human brain microvascular endothelial cells (HBMVECs) were encapsulated in an ECM-based hydrogel used to embed the COs. During the expansion and maturation phases, organoid media was supplemented with endothelial cell growth (ECG) media and VEGF. By adopting this approach, we generated vascularized brain organoids with key morphological and physiological features of the cerebrovasculature, establishing a promising physiologically relevant model for a wide range of research applications.

**Figure 1.**
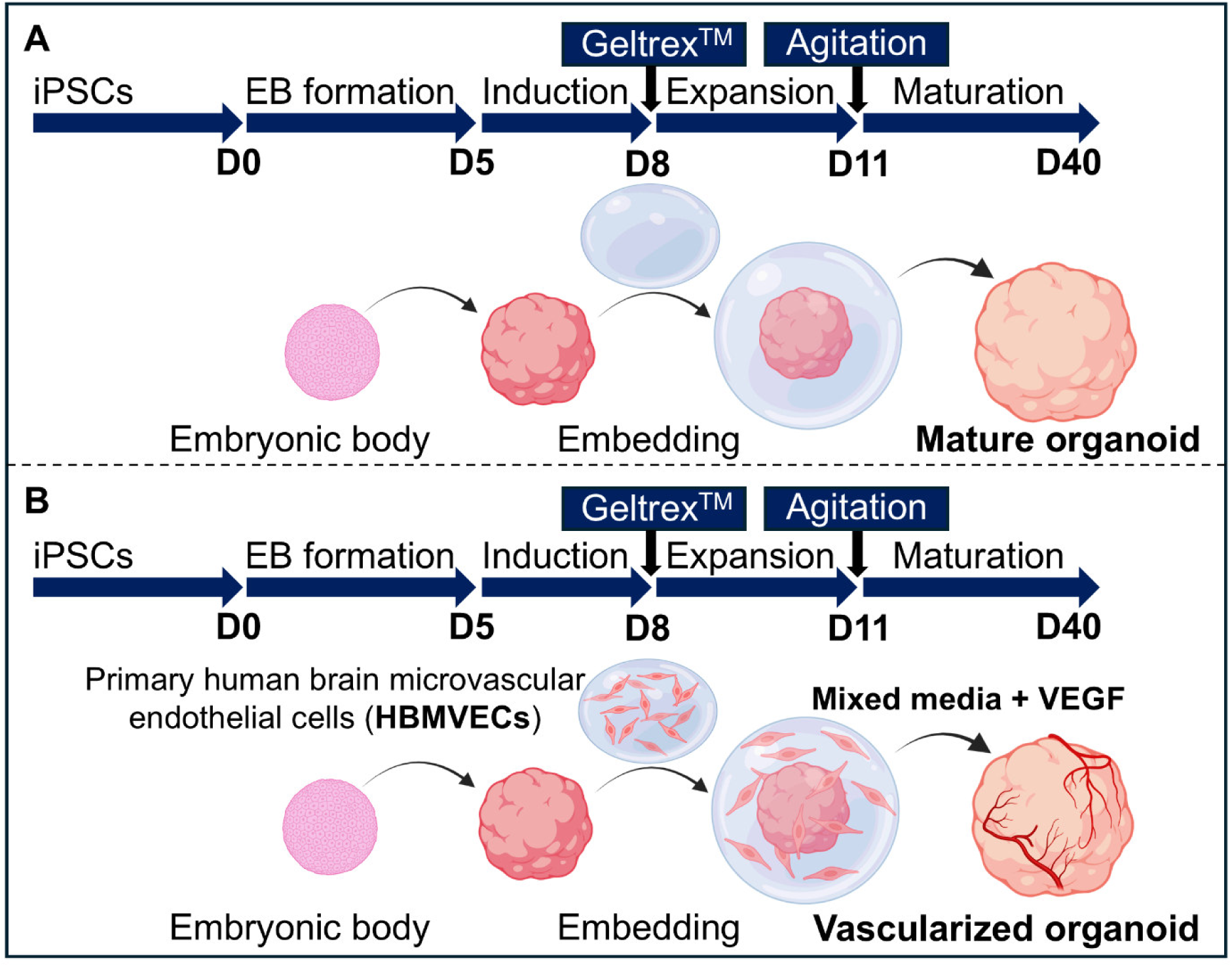
Schematic overview of the brain organoid vascularization strategy. (A) Graphical representation of the cerebral organoid depicting starting induced pluripotent stem cells (iPSCs, Day 0), embryonic formation (D0-D5), induction of the neuroepithelium (D5-D8), expansion of the neural tissue (D8-D11) and maturation of the organoid (D11-D40+). (B) Graphical representation of the protocol for generating vascular-like networks in the cerebral organoid. Images were created using Biorender.com.

## RESULTS

### HBMVECs network formation is modulated by the hydrogel concentration, media composition and VEGF dosage

HBMVECs were chosen for incorporation in the COs because of their high specialized phenotype and tight junction protein expression^50^. To determine the most effective incorporation strategy, two different procedures were tested: *i)* addition of HBMVECs for their attachment after Geltrex^TM^ polymerization (surface attachment approach) and *ii)* encapsulation of HBMVECs within the Geltrex^TM^ matrix (encapsulation approach) (Figure S1A). Although endothelial networks were identified on day 9 using both approaches, those formed on the surface of the droplet deteriorated faster over time than the more stable encapsulated networks up to day 11 (Figure S1B). Thus, it was determined that the encapsulation of ECs within the matrix was the best approach to control cell density, spatial distribution, and network stability. By comparing COs from the same A-iPSC line grown with different encapsulating cell densities, we determined that 50,000 HBMVECs per organoid (2,000 cells per µL of Geltrex^TM^) was optimal for forming superficial networks while preventing an excess of ECs from depositing in layers onto the organoid surface, as observed at higher cell numbers (500,000 and 2 million cells) (Figure S1C and D).

To optimize HBMVECs tube formation, we developed an assay to investigate cell behavior when encapsulated in Geltrex^TM^ for up to 10 days (Figure 2A). First, we evaluated how hydrogel concentration affected endothelial network connectivity and the role of supplementing ECG media with 50 ng/mL of VEGF^36^. As expected, hydrogel percentage strongly influenced endothelial networks assembly, with lower concentrations providing a more tunable matrix for HBMVECs. For instance, we identified the highest network density in 40% Geltrex^TM^ with and without VEGF, exhibited the highest percentage of network area, greatest total vessel length and lowest lacunarity value (Figure 2B, C). In contrast, endothelial connections were barely formed at the highest hydrogel concentration tested (80% Geltrex^TM^) (Figure 2B). VEGF supplementation significantly increased the percentage of vessel area and total vessel length while decreasing the lacunarity values, indicating more interconnected endothelial networks. In addition, the VEGF effect was more pronounced at 40% and 60% concentrations (Figure 2C); thus, we hypothesized that reducing the Geltrex^TM^ concentration below the standard 100% used in CO protocols, combined with VEGF supplementation, would enhance network formation by encapsulated HBMVECs,

**Figure 2.**
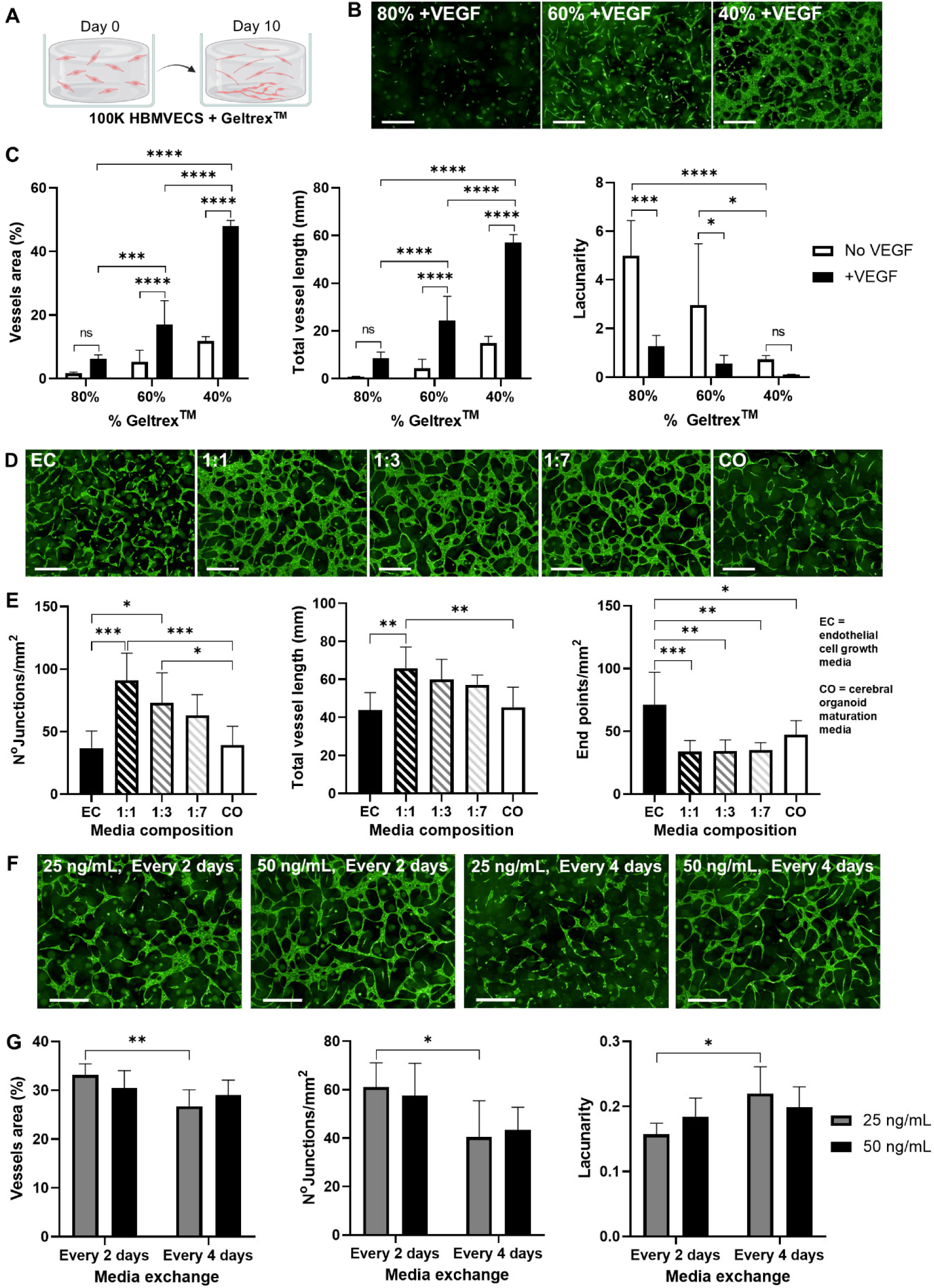
Vessel formation with human brain microvascular endothelial cells (HBMVECs) is enhanced at lower Geltrex^TM^ concentrations and by mixed endothelial-organoid media in the presence of vascular endothelial growth factor (VEGF) (A) Graphical representation of 100,000 HBMVECs encapsulated in different Geltrex^TM^ concentrations and grown for 10 days using different media and VEGF doses. Created with Biorender.com. (B) Representative confocal images of Calcein AM-stained endothelial networks formed in 80%, 60%, and 40% Geltrex^TM^ supplemented with 50 ng/mL VEGF for 10 days. Scale bars: 500 µm. (C) Percentage of vessel area (%), total vessel length (mm) and lacunarity values of endothelial networks formed in 80%, 60%, and 40% Geltrex^TM^, with or without 50 ng/mL VEGF. Mean ± SD, N = 6 independent wells, two repeats. Two-way ANOVA; asterisks showing Tukey’s post hoc comparisons; ns = not significant (p > 0.05); *p < 0.05, ***p < 0.001, ****p < 0.0001. (D) Representative confocal images of Calcein AM-stained endothelial networks grown in 50% Geltrex^TM^ with different media (endothelial cell growth media (ECG), organoid maturation media (CO) or mixed media at 1:1, 1:3 and 1:7 EC:CO ratios), all supplemented with 50 ng/mL VEGF for 10 days. Scale bars: 500 µm. (E) Total vessel length (mm), number of junctions per mm^2^, and end points per mm^2^ of endothelial networks grown in different media compositions. Mean ± SD, N = 6 independent wells, two repeats. One-way ANOVA; asterisks showing Tukey’s post hoc comparisons; *p < 0.05, **p < 0.01, ***p < 0.001. (F) Representative confocal images of Calcein AM-stained endothelial networks formed in 50% Geltrex^TM^ with 1:7 mixed media, supplemented with 25 or 50 ng/mL VEGF, with media exchanges every 2 or 4 days for 10 days. Scale bars: 500 µm. (G) Percentage of vessel area (%), number of junctions per mm^2^, and lacunarity values of endothelial networks formed at different VEGF concentrations (25, 50 ng/mL) and media exchange frequency (every 2 or 4 days). Mean ± SD, N = 6 independent wells, two repeats. Two-way ANOVA; asterisks showing Tukey’s post hoc comparisons; *p < 0.05.

To simultaneously balance neural tissue expansion while stimulating vasculogenesis and angiogenesis, we investigated the influence of organoid maturation media HBMVEC vessel formation. ECG media was mixed with maturation media at increasing ratios (1:0, 1:7, 1:3, 1:1 and 0:1) and supplemented with 50 ng/mL VEGF. All mixed media conditions (1:1, 1:3 and 1:7) promoted better network assembly than ECG or maturation media alone, with mixed media 1:1 forming the most robust networks (highest total vessel length, highest number of junctions and lowest number of end points) (Figure 2D, E). However, the lowest ECG media ratio (1:7) was selected for further experiments, to minimize the impact on CO development.

To align VEGF treatment with the organoid media change schedule (every 4 days), we assessed the effect of two doses (25 and 50 ng/mL) every 2 or 4 days. Network analysis showed that 25 ng/mL VEGF every 4 days resulted in poorer network formation (Figure 2F, G). Therefore, since no significant differences were observed with the 50 ng/mL dose, it was selected for the co-cultures.

### Integration of HBMVECs into cerebral organoids generates more consistently shaped microtissues without impacting stiffness

Cerebral organoids were successfully developed from two quality-checked iPSC lines (A-iPSCs and S-iPSCs, see Experimental model and study participant details and Figure S2) using the STEMdiff™ Cerebral Organoid Kit. COs presented typical features such as SOX2+ neuroepithelia/neural rosettes, PAX6+ layers in the cortical-like regions and βIIITUB+ neurons in the most mature areas (Figure S3A-C). To generate vascularized COs, HBMVECs were encapsulated within a 80% Geltrex^TM^ droplet (+BECs), and the organoids were cultured in the previously optimized 1:7 ECG:CO mixed media (hereinafter just mixed media or MM) supplemented with 50 ng/mL VEGF (+VEGF) (Figure 1). Control COs were grown in media provided by the kit without encapsulating HBMVECS (CTRL). Overall, six testing conditions were evaluated to assess the impact of each protocol adaptation: three without encapsulating HBMVECS (CTRL, +VEGF, MM+VEGF) and three with encapsulating HBMVECS (BECs CTRL, BECs+VEGF, BECs MM+VEGF) (Figure 3).

**Figure 3.**
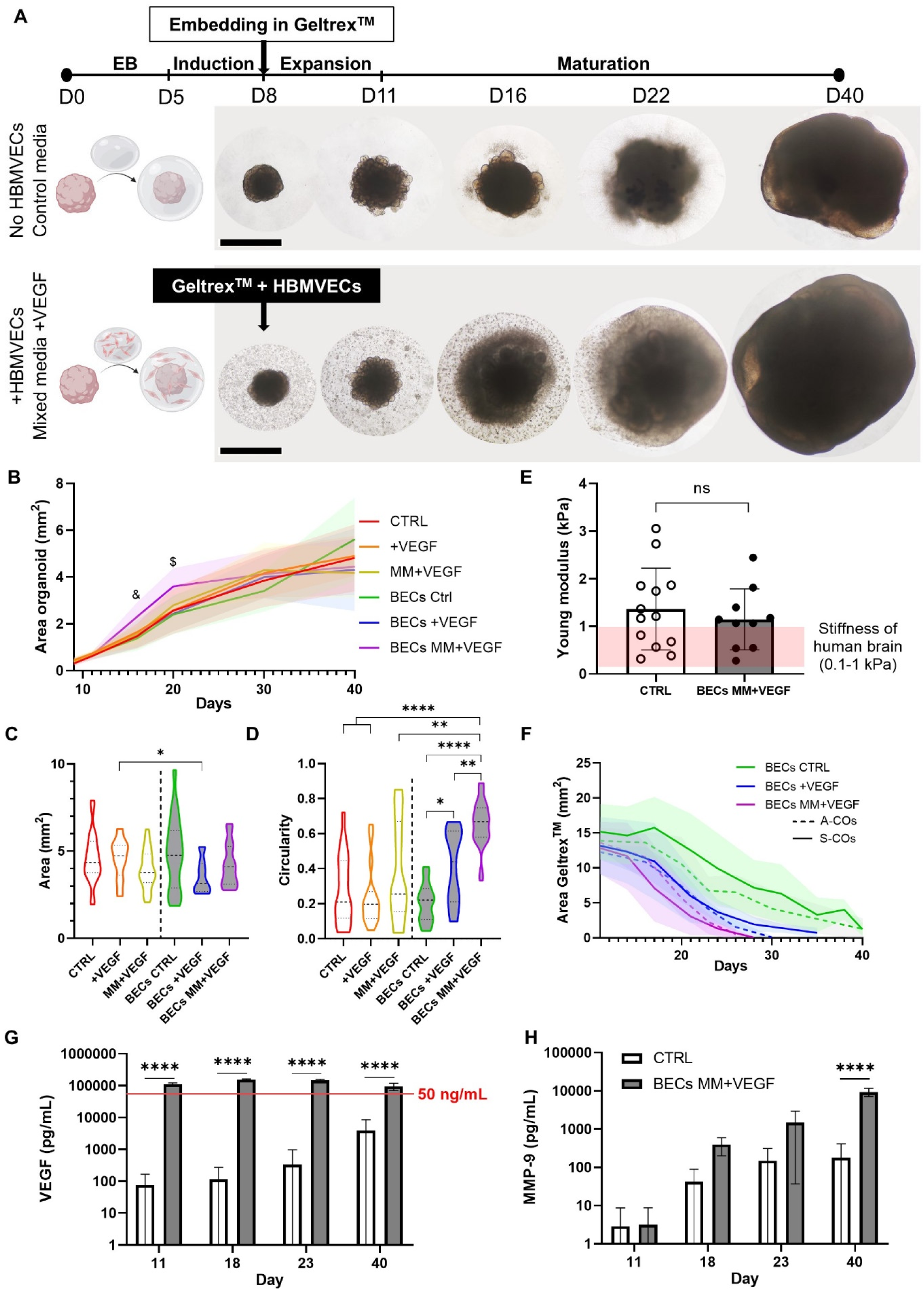
Promoting vascularization of the organoid accelerates early organoid growth and spherical morphology without impacting on the size or tissue stiffness by day 40. (A) Brightfield images of standard organoids grown according to the Lancaster protocol^9^ (CTRL) compared to organoids grown in the modified protocol, where HBMVECs were encapsulated in a Geltrex^TM^ droplet and grown in mixed media supplemented with VEGF (BECs MM+VEGF). Representative images were captured on days 8, 11, 16, 22 and 40. Scale bar: 1 mm. (B) Organoid size evolution from encapsulation until day 40 across the six tested conditions: control (red), +VEGF (orange), MM+VEGF (yellow), BECs CTRL (green), BECs +VEGF (dark blue), and BECs MM+VEGF (purple). Organoid size was measured as the area in the brightfield images (mm^2^). Mean ± SD, N = 16-25, five independent batches, two different iPSC lines. Mixed-effect model, followed by Tukey’s post hoc comparisons; on day 16, BECs MM+VEGF were significantly different (&) from CTRL (p = 0.001), MM+VEGF (p = 0.0013), BECs CTRL (p = 0.0002) and BECs +VEGF (p = 0.0085). On day 20, BECs MM+VEGF were significantly different ($) from CTRL (p = 0.0143), +VEGF (p = 0.0128), MM+VEGF (p = 0.0027), BECs CTRL (p = 0.0016) and BECs +VEGF (p = 0.0007). (C) Largest longitudinal area (mm^2^) of day 40 organoids across all conditions (white-filled plots: absence of HBMVECs; grey-filled plots: encapsulated HBMVECs). The violin shape represents the data distribution, median (thick line), interquartile range (thin lines), N = 19-31, 5 independent batches, two different iPSC lines. Brown-Forsythe and Welch ANOVA test, asterisks showing Games-Howell’s post hoc comparisons; *p < 0.05. (D) Organoid circularity at day 40 across all conditions (white-filled plots: absence of HBMVECs; grey-filled plots: encapsulated HBMVECs). The violin shape represents the data distribution, median (thick line), interquartile range (thin lines), N = 14-27, five independent batches, two different iPSC lines. Brown-Forsythe and Welch ANOVA test, asterisks showing Games-Howell’s post hoc comparisons; *p < 0.05, **p < 0.01, ****p < 0.0001. (E) Nanoindentation measurements of tissue stiffness (Young modulus, in KPa) of CTRL and BECs MM+VEGF organoids. Mean ± SD, N = 10-13, two independent batches, two different iPSC lines. Unpaired T-test; ns = not significant (p > 0.05). (F) Degradation of the Geltrex^TM^ droplet area (mm^2^) over time (white-filled plots: absence of HBMVECs; grey-filled plots: encapsulated HBMVECs; dash lines represent organoids generated with the A-iPSC line; uninterrupted lines represent organoids generated with the S-iPSC line). Mean ± SD, N = 9-15, two or three independent batches per each of the two IPSC lines. (G) ELISA quantification of VEGF in culture supernatants from CTRL and BECS MM+VEGF organoids. The red line represents the 50 ng/mL VEGF dose added in the MM+VEGF media. Mean ± SD, N = 4, two independent batches for each of the two iPSC lines. Two-way ANOVA, asterisks showing Sidak’s post hoc comparisons; ****p < 0.0001. (H) ELISA quantification of matrix metalloproteinase 9 (MMP-9) in culture supernatants from CTRL and BECS MM+VEGF organoids. Mean ± SD, N = 4, two independent batches for each of the two iPSC lines. Two-way ANOVA, asterisks showing Sidak’s post hoc comparisons; ****p < 0.0001.

All the COs had uniform sizes at the encapsulation step, and from there, the size was monitored until culture day 40 (Figure 3A). In all tested conditions, COs developed neuroepithelial budding between days 8 and 16, as shown in the brightfield images (Figure 3A). However, BECs MM+VEGF COs grew significantly faster than organoids in the other conditions up to day 22 (Figure 3B). Morphological analysis at the final timepoint (day 40) showed no differences in size across conditions, except for BEC+VEGF organoids, which were significantly smaller than their controls without HBMVECs (+VEGF, Figure 3C). Remarkably, BECs+VEGF and BECs MM+VEGF organoids were more uniform in size than BECs CTRL organoids, across batches. Size uniformity correlated with significantly higher circularity values for BECs+VEGF and BECs MM+VEGF organoids compared to BECs CTRL (Figure 3D). In contrast, varying media conditions had no effect on the circularity of organoids without HBMVECs. We observed that some of these organoids retained traces of non-invaded Geltrex^TM^ or exhibited irregular neurites projecting in multiple directions, which contributed to the lower circularity. There was no difference in these findings between the two iPSC lines. To assess whether smaller, rounder organoids exhibited denser cerebral tissue, we performed a nanoindentation assessment to compare the compression stiffness of batch-matched CTRL and BECs MM+VEGF organoids (Figure 3E). In comparison with the physiological stiffness of the brain tissue (0.1-1 kPa)^51^, CO stiffness ranged between 0.5-3 kPa, with no significant difference between the two organoid groups for either of the iPSC lines. This indicates that the external pressure exerted by the surrounding endothelial network remodeling did not affect the whole tissue rigidity.

Differences in the Geltrex^TM^ droplet degradation rates were observed among the +BECs conditions, by tracking the area of free Geltrex^TM^ (excluding the area occupied by the organoid) over 40 days. Compared to the CTRL media, Geltrex^TM^ droplets containing HBMVECs appeared to shrink faster when cultured in MM+VEGF media, followed by +VEGF media. We hypothesize that pro-angiogenic ECs exhibit enhanced efficacy degrading the surrounding matrix (Figure 3F), as VEGF promotes ECM remodeling and angiogenesis by inducing strong expression of matrix metalloproteinases, such as MMP-9, in ECs^52–54^. To confirm this, culture supernatants of BEC MM+VEGF and CTRL organoids were collected and the concentrations of VEGF and MMP-9 were quantified. Fresh media were supplemented with 50 ng/mL VEGF; however, after 3-4 days of culture, we found that VEGF levels were sustained at over 100 ng/mL (Figure 3G), indicating a positive feedback loop in which ECs released more VEGF. In contrast, CTRL organoids revealed basal and increasing VEGF secretion over time reaching 4 ng/mL on day 40, probably due to the growing hypoxic core. Simultaneously, MMP-9 levels were also higher in BECs MM+VEGF supernatants than in the CTRL, reaching significance on day 40 (Figure 3H). MMP-9 expression in BECs MM+VEGF increased exponentially to almost 10 ng/mL on day 40, while in the CTRL cultures MMP-9 secretion appeared to plateau between 150-200 pg/mL between the two later time points (day 23 and day 40).

### HBMVECs form multicellular networks on the organoid surface that can be modulated by the Geltrex^TM^ concentration

To reveal the extent of endothelial cells on the organoids surface at day 40, whole organoids were immunostained for ECs (CD31+) and neurons (βIIITUB+, Figure 4A). In this case, notable differences were noticed between the two iPSC lines tested. For the A-iPSC-generated cerebral organoids (A-COs), endothelial networks formed regardless of HBMVECs addition or media composition (Figure 4A top panel), with high variability in network coverage even among organoids from the same batch (Figure 4B). In contrast, cerebral organoids generated with the S-iPSCs line (S-COs) were mostly devoid of spontaneously generated ECs in the same HBMVECs-absent conditions, except for some noticeable in the MM+VEGF group (Figure 4A top panel, C). As predicted, the formation of endothelial networks in the HBMVEC-containing organoids was enhanced by the media supplementation with VEGF (Figure 4A middle panel, Figure 4B and C). BECs CTRL organoids exhibited around 5% CD31+ surface area, increasing to 10-15% with VEGF in both A– and S-COs. In BECs MM+VEGF organoids, A-COs exhibited a significant increase in endothelial networks reflected by a higher superficial CD31+ area, which was not consistently observed for the S-COs. Morphological analysis of these vascular-like constructs revealed that the BECs MM+VEGF condition resulted in a lower number of vessel end points, increased average vessel length and higher density of junctions when compared to BECs CTRL (Figure 4D). Altogether, BECs MM+VEGF organoids generated multicellular and highly interconnected superficial endothelial networks, which were abundant in A-COs, but less prevalent in S-COs (Figure 4A, bottom panel). Some of these networks exhibited hierarchical organization, with smaller branches extending from wider vessels (Figure S4).

**Figure 4.**
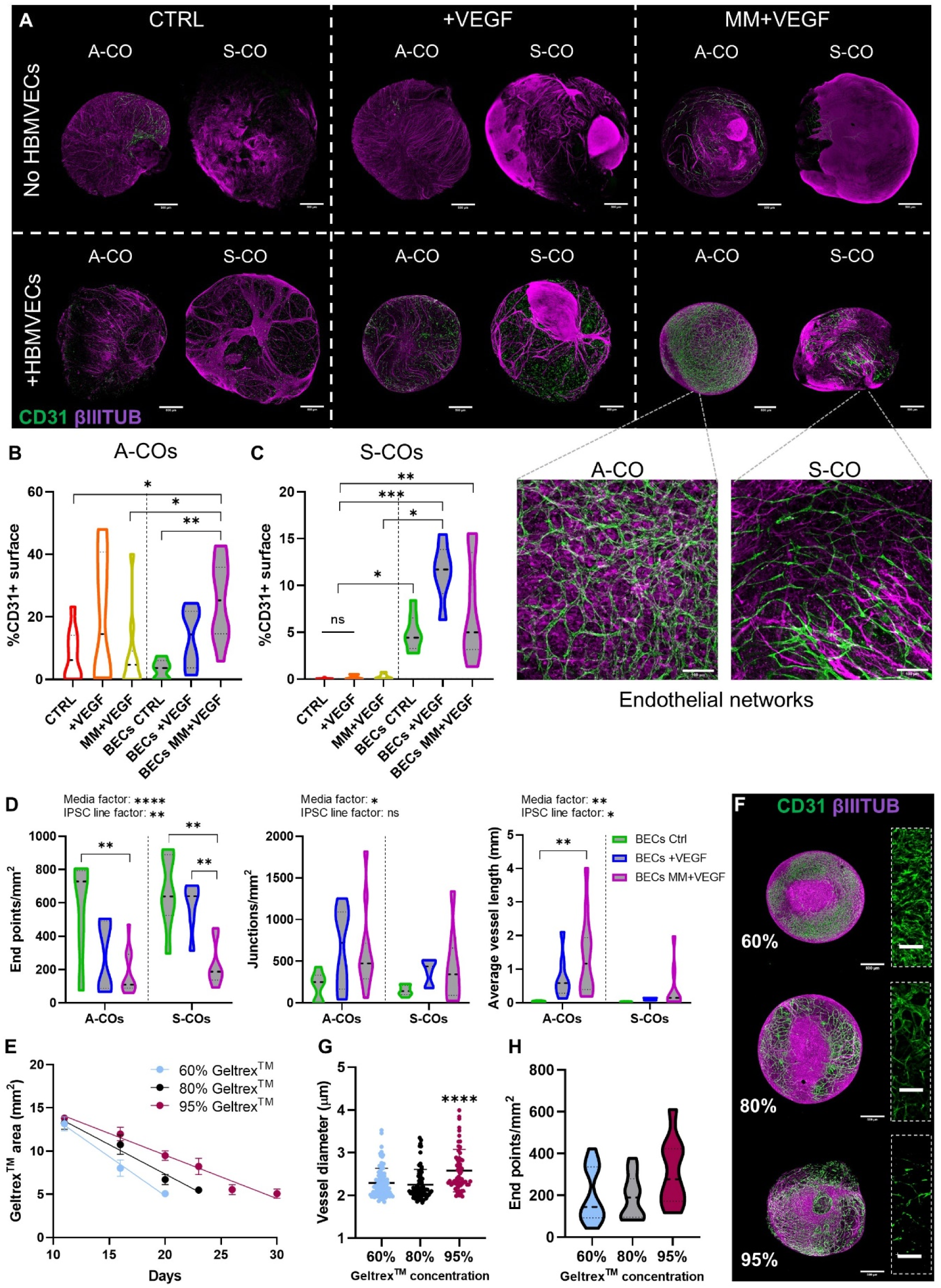
Abundant multicellular endothelial networks are formed on the surface of cerebral organoids, with morphological features dependent on media conditions or Geltrex^TM^ concentration. (A) Confocal images of A and S-COs grown under tested vascularization conditions, wholemount stained for CD31 (green) and βIIITUB (magenta). Dashed lines separate the different conditions: media used (control, +VEGF or MM+VEGF, vertical display), presence or absence of encapsulated HBMVECs (horizontal display). Within each condition, a comparison between organoids generated with the A-and S-iPSCs lines. Scale bars: 500 µm. At the bottom, higher magnification confocal images showing endothelial networks on the surface of BECs MM+VEGF organoids, for both iPSCs lines. Scale bars: 100 µm. (B) Percentage of CD31+ surface in A-COs. Control (red), +VEGF (orange), MM+VEGF (yellow), +BECs CTRL (green), +BECs +VEGF (dark blue) and +BECs MM+VEGF (purple); no HBMVECs, white filling; +HBMVECs, grey filling. Violin shape represents data distribution, Median (thick line), interquartile range (thin lines), N = 6-12, three independent batches. Brown-Forsythe and Welch ANOVA test, asterisks showing Games-Howell’s post hoc comparisons; *p < 0.05, **p < 0.01. (C) Percentage of CD31+ surface in S-COs. Same color system than above. The violin shape represents data distribution, median (thick line), interquartile range (thin lines), N = 6-8, two independent batches. Kruskall-Wallis test, asterisks showing Dunn’s post hoc comparisons; *p < 0.05, **p < 0.01, ***p < 0.001. (D) Quantification of network end points per mm^2^, average vessel length (mm), and junctions per mm^2^ of the HBMVECs-containing organoids, separated by iPSCs origin. BECs CTRL (green), BECs +VEGF (dark blue) and BECs MM+VEGF (purple). The violin shape represents the data distribution, median (thick line), interquartile range (thin lines), N = 5-12, three or two independent batches for each line. Two-way ANOVA test; media and iPSC line factor significance expressed on top of each graph. Tukey’s post hoc comparisons are represented by brackets over relevant bars; ns = not significant (p > 0.05), *p < 0.05, **p < 0.01, ***p < 0.001, ****p < 0.0001. (E) Degradation rate of different concentrations of Geltrex^TM^ droplets in BECs MM+VEGF conditions, measured by the area change (mm^2^) over time. Light blue (60%), black (80%), Bordeaux (95%). Mean ± SD, N = 5, one batch of organoids. (F) Representative confocal images of BECs MM+VEGF A-COs grown at different Geltrex^TM^ concentrations (60%, 80% and 95%), wholemount stained for CD31 (green) and βIIITUB (magenta). Scale bars: 500 µm. On the right, higher magnification images of the endothelial networks are shown. Scale bars: 100 µm. (G) Vessel diameter (µm) of the superficial endothelial networks formed under different Geltrex^TM^ concentrations. Light blue (60%), black (80%), Bordeaux (95%). Mean ± SD, N = 9-22 (individual organoids) and >5 areas of interest per organoid analyzed, five different batches, same A-iPSC line. Kruskal-Wallis test, asterisks showing Dunn’s post hoc comparisons; ****p < 0.0001. (H) End points per mm^2^ of the superficial endothelial networks formed under different Geltrex^TM^ concentrations. Light blue (60%), white (80%), and Bordeaux (95%) colors were used. The violin shape represents the data distribution, median (thick line), interquartile range (thin lines), N = 9-22, five different batches, and the same A-iPSC line. One-way ANOVA test, Tukey’s post hoc comparisons; not significant (p > 0.05).

Using A-COs, we investigated the role of Geltrex^TM^ concentration in the formation of these vascular-like networks. BECs MM+VEGF COs were encapsulated in increasing concentrations of Geltrex^TM^ (60, 80 and 95%) with 50,000 HBMVECs. As expected, the rate of Geltrex^TM^ degradation (calculated by monitoring the droplet area over time) was inversely proportional to its concentration (Figure 4E). This degradation rate influenced the speed at which ECs reached the organoid surface, ultimately shaping endothelial network morphology (Figure 4F). Notably, vessel diameters were significantly wider in the 95% Geltrex^TM^ condition compared to the 80% and 60% conditions (Figure 4G). Morphological analysis revealed that networks in the 60% condition were generally more branched and interconnected (larger vessels area, higher junction density, longer average vessel length and lower number of vessels end points) whereas networks in the 95% condition appeared more spaced and discontinuous (Figure 4H and S5).

### Endothelial networks in COs are a mix of HBMVECs and endogenously differentiated ECs

To investigate the source of endogenous EC differentiation in A-COs, we examined the presence of mesodermal progenitors in early COs on days 5 and 8. Immunostaining for the mesodermal progenitor marker brachyury was absent on the surface of early organoids (Figure 5A). In contrast, day 8 COs exhibited large OTX+ areas, an ectodermal progenitor marker, which appeared fainter in areas with visible neural rosettes (Figure 5B), consistent with what was previously reported^55^. These findings suggest the absence of mesodermal progenitors in the iPSCs cultures and confirm successful neuroectodermal induction.

**Figure 5.**
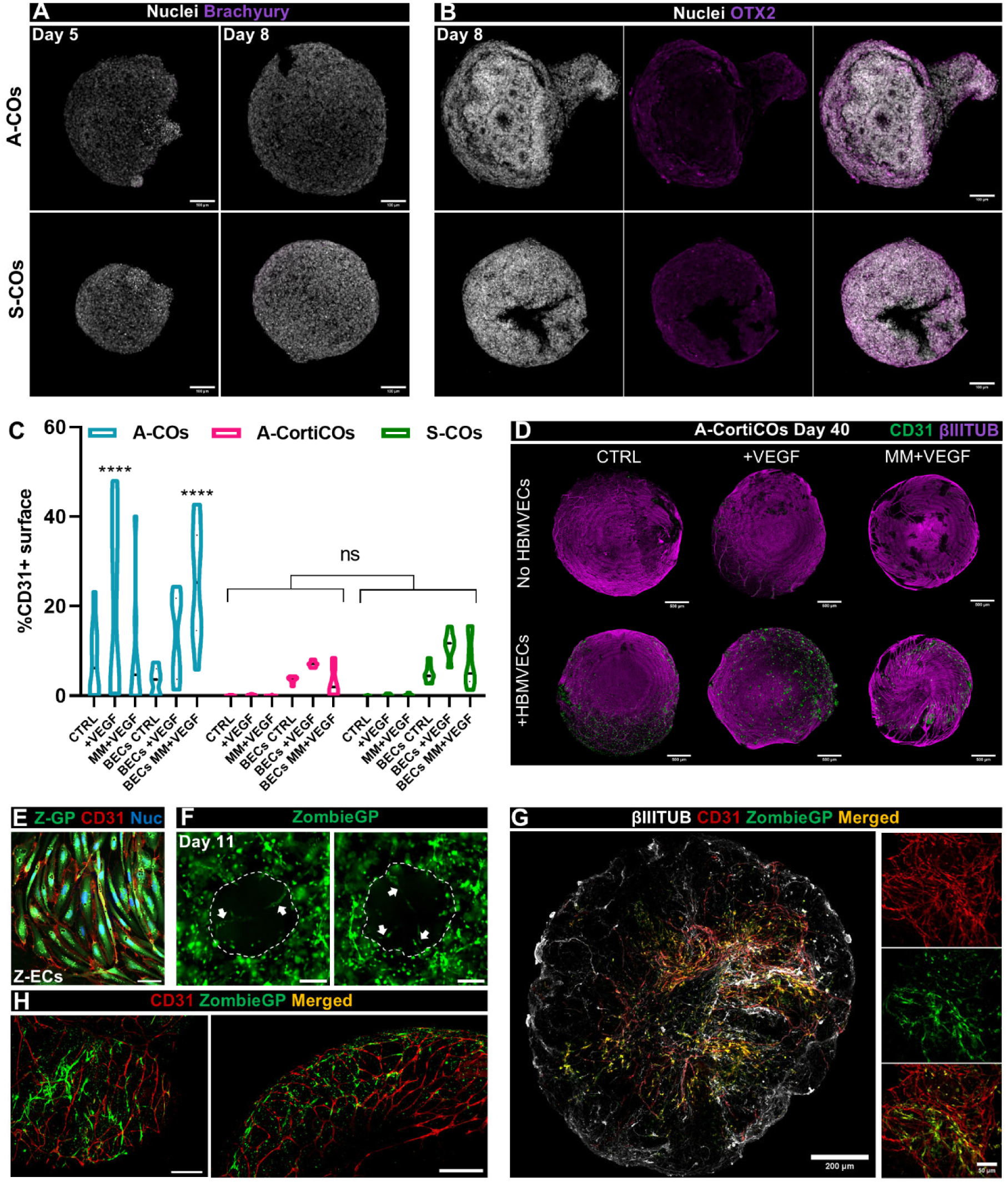
Endothelial networks formed in COs are partially formed by endogenously differentiated ECs, which can be repressed by dual SMAD inhibition. (A) Confocal images of day 5 embryonic bodies and day 8 early cerebral organoids stained for brachyury (purple) and nuclei (white). Top, A-COs; bottom, S-COs. Scale bars: 100 µm. (B) Confocal images of day 8 early cerebral organoids stained for OTX2 (purple) and nuclei (white). Top, A-COs; bottom, S-COs. Scale bars: 100 µm. (C) Percentage of CD31+ coverage on day 40 A-COs (blue), A-CortiCOs (pink), and S-COs (green) grown under identical vascularization conditions. The violin shape represents the data distribution, median (thick line), N = 6-12, two or three different rounds per organoid type. Two-way ANOVA, asterisks showing Tukey’s post hoc comparisons; ns = not significant (p > 0.05), *p < 0.05, and ****p < 0.0001. (D) Confocal images of Day 40 A-CortiCOs whole stained for CD31 (green) and βIIITUB (purple), grown under different vascularization conditions: CTRL, +VEGF, MM+VEGF, BECs CTRL, BECs +VEGF, and BECs MM+VEGF. Scale bars: 500 µm. (E) Confocal image of fluorescent iPSC-derived endothelial cells (Z-ECs) expressing Zombie Green protein (Z-GP) in the cytoplasm, grown after the magnetic sorting, and stained for CD31 (red) and nuclei (blue). Scale bar: 200 µm. (F) Fluorescence microscopy image of day 11 COs embedded in a droplet of 80% Geltrex^TM^ encapsulating Z-ECs. White arrows indicate endothelial tubes penetrating the developing neural tissue. Scale bars: 200 µm. (G) Panel of confocal microscopy images revealing a tissue-cleared cerebral organoid with dual-origin endothelial cell types interacting to form networks: endothelial cells (CD31, red), Z-ECs added to the organoid (Zombie Green protein, green), co-localization (yellow), and βIIITUB (white). Scale bars: 200 µm (left) and 50 µm (right). (H) Confocal images of day 40 cerebral organoids encapsulating Z-ECs (green) stained for CD31 (red), showing areas in which the green fluorescent cells were not CD31+. Scale bars: 200 µm.

Next, we generated cortical organoids (CortiCOs) with the A-iPSC line using a dual SMAD signaling inhibition protocol, which strongly represses mesodermal differentiation^17^. CortiCOs displayed SOX2+ neuroepithelial areas and βIIITUB+ differentiated areas, similar to the COs (Figure S3). When the same vascularization conditions as those for the COs were applied to A-CortiCOs, there was almost no presence of ECs in the HBMVEC-absent conditions, especially when compared to A-COs (Figure 5C and D, and 4A). In fact, CD31+ coverage values across conditions were comparable to those previously reported for S-COs (Figure 4A). While the CD31+ surface area was similar among all BECs CTRL and BECs +VEGF organoids, A-COs showed a significantly higher percentage of superficial networks than A-CortiCOs and S-COs, which displayed similar values (Figure 5C).

To trace endothelial cell development and network formation in COs, fluorescent iPSC-derived ECs (Z-ECs) were used to encapsulate MM+VEGF organoids instead of HBMVECs. Z-ECs were obtained by differentiation from a Zombie Green protein (GP)-expressing iPSC line and characterized for the endothelial phenotype (Figure 5E and S6). Leveraging this fluorescent labelling, we monitored the invasion of endothelial networks into and around the organoids (Figure 5F). At day 40, whole organoids were stained for CD31 and βIIITUB and tissue cleared, revealing that vascular-like networks in both A– and S-COs were composed of both endogenous, CO-derived ECs (CD31+) and exogenously added Z-ECs (ZombieGP and CD31+) (Figure 5G). Thus, we confirmed the dual origin of these endothelial networks, including those found in S-COs. We noticed that some Z-ECs were not CD31+ (Figure 5H), potentially due to either Z-ECs transdifferentiation into other cell types or insufficient CD31 antibody penetration into the tissue regions where these fluorescent Z-ECs were detected.

### Vascular-like networks are present within the organoid and exhibit BBB characteristics

This study aimed to determine whether the vascularization method effectively established networks capable of penetrating neural tissue and integrating within the organoids. Immunostaining of cryosections revealed that CD31+ network-organized endothelial cells were present within the BECs MM+VEGF COs (herein ‘vascularized’) in a significantly higher number than batch-matched control COs (Figure 6A, B). These endothelial cells were mainly found in the lower density cell areas of the organoids (normally at the edges) and did not penetrate the neuroepithelial areas. Endothelial tubes also expressed adhesion molecule vascular endothelial cadherin (VE-CAD) (Figure 6C). As expected, there were fewer internalized endothelial networks in the S-COs (Figure 6D).

**Figure 6.**
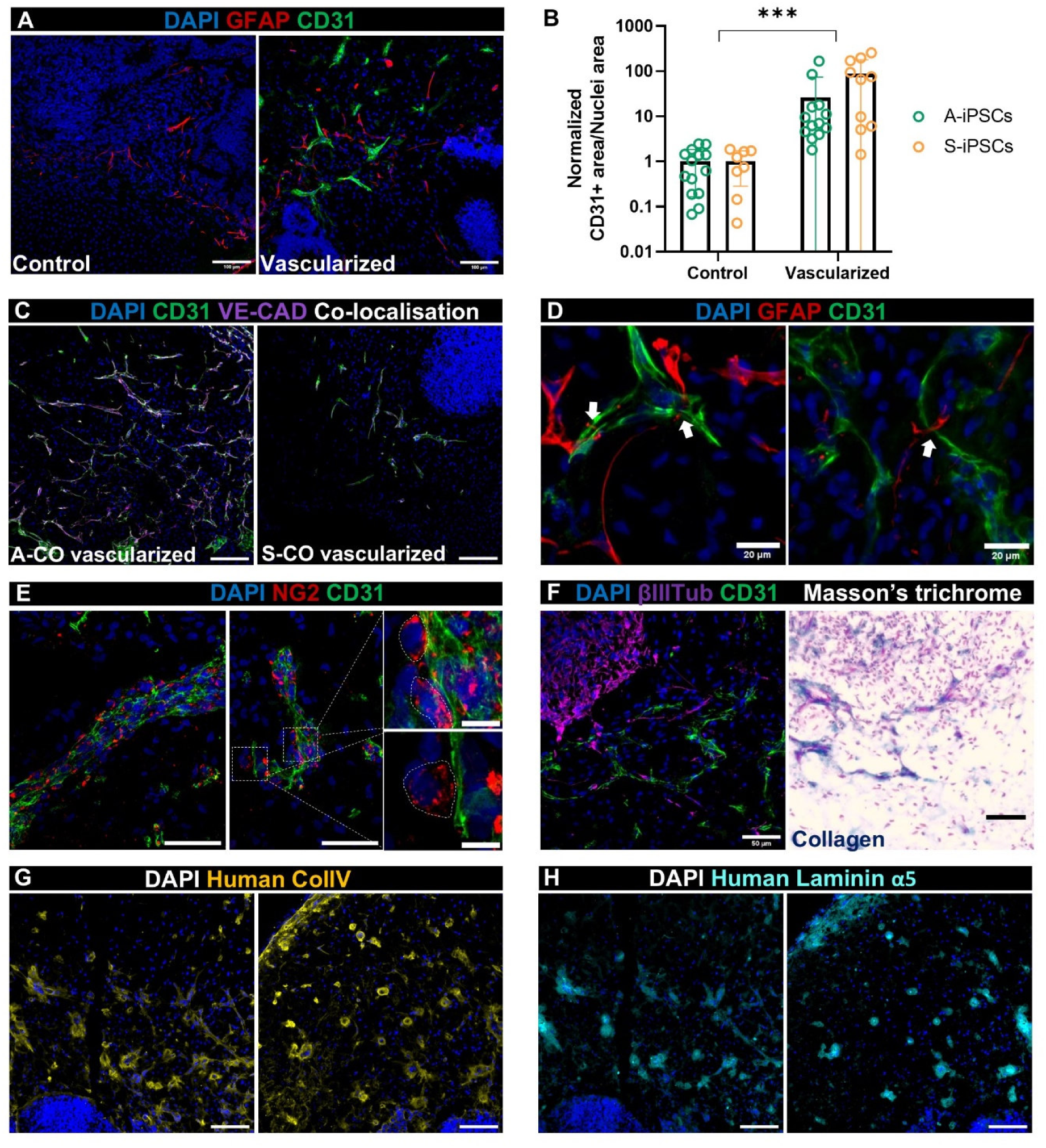
Vascularized COs display characteristics of the blood-brain barrier, such as astrocytes wrapping the endothelial tubes, high expression of VE-CAD, and collagen IV-laminin depositions surrounding the networks. (A) Representative confocal images of middle cryosections of a control and vascularized CO stained for CD31 (green), GFAP (red) and nuclei (blue). Scale bars: 100 µm. (B) CD31+ area/nuclei area in control versus vascularized COs sections. The values were normalized to the average of the batch-matched controls. Mean ± SD, N = 8-14, three or four batches for each iPSC line. Two-way ANOVA; vascularization effect: ***p < 0.001. (C) Representative confocal images of vascularized COs generated from both A-iPSC and S-iPSC lines, stained for CD31 (green) and VE-CAD (purple), showing their co-localization in the intercellular junctions of the networks (white). Scale bars: 100 µm. (D) Confocal images showing the interaction (white arrows) between astrocytes (GFAP+, red) and endothelial tubes (CD31+, green). Scale bars: 20 µm. (E) Confocal images of neural/glial antigen 2 (NG2, red) and CD31 (green) immunostaining of vascularized COs sections, with high magnification images showing individual pericyte-like cells attached to endothelial networks (white line). Scale bars: 50 µm (left and centre) and 10 µm (right). (F) Comparison of the same cerebral organoid area immunostained for βIIITUB (neurons, purple) and CD31 (green) on the left, and histological staining with Masson’s trichrome on the right (cells in pink, collagen in dark blue). Scale bars: 50 µm. (G) Representative confocal images of vascularized COs regions highly populated with endothelial networks stained for human collagen IV (yellow) and nuclei (blue). Scale bars: 100 µm. (H) Same areas as in (F) but stained for human-specific laminin α5 (cyan) and nuclei (blue). Scale bars: 100 µm.

Co-staining with the astrocytic marker glial fibrillary acidic protein (GFAP) showed that our adapted CO protocol did not inhibit astrocyte formation (Figure 6A). Astrocytes were identified in direct contact with the endothelial tubes, as typically encountered in the neurovascular unit (Figure 6D), and abundant neural/glial antigen 2 (NG2)-positive cells were detected attached in the proximity of endothelial networks in a pericyte-like fashion (Figure 6E). In addition, collagen deposition (dark blue) surrounding the CD31+ networks was revealed by Masson’s trichrome staining (Figure 6F). We confirmed using human-specific antibodies that these depositions around the endothelial structures in vascularized COs were mainly composed of collagen type IV and laminin α5 (Figure 6G, H).

### Vascularized COs are more permeable to nutrients and displayed smaller apoptotic areas

Vascularization of organoids introduced significant morphological changes; however its physiological implications remain unclear. To investigate whether the vascular-like networks were lumenized and perfusable, we incubated control and vascularized COs in a fluorescent 70 kDa dextran-rhodamine B (d-RhoB) solution, transferred them to PBS, and quantified the d-RhoB that was held in the tissue (Figure S7A). Fluorescence levels in the vascularized COs were significantly higher than those in the control COs (Figure S7B). Because this effect could be influenced by organoid size, we plotted the fluorescence intensity against organoid size revealing two significantly different linear regressions between the two groups. At smaller sizes, vascularized COs transferred more d-RhoB than size-matched control COs, whereas fluorescence in the controls was size-independent (Figure S7C). Despite these promising results, we could not determine whether d-RhoB was retained within the networks or whether there were open lumens. For further insights, after d-RhoB incubation, organoids were cryopreserved and sectioned and examined for d-RhoB internalization. As expected, d-RhoB mainly penetrated the periphery of the tissue (Figure 7A). However, regions with higher fluorescence intensity coincided with the densest endothelial areas in vascularized A-COs. Quantification of the d-RhoB+ area across the middle cryosections indicated that vascularized A-COs were significantly more penetrable than control A-COs, whereas no differences were observed in S-COs (Figure 7B). Furthermore, noticeably stronger d-RhoB+ halos were found alongside some endothelial networks (Figure 7C), together with the presence of open lumens as confirmed by hematoxylin and eosin (H&E) staining of paraffin-embedded sections (Figure 7D).

**Figure 7.**
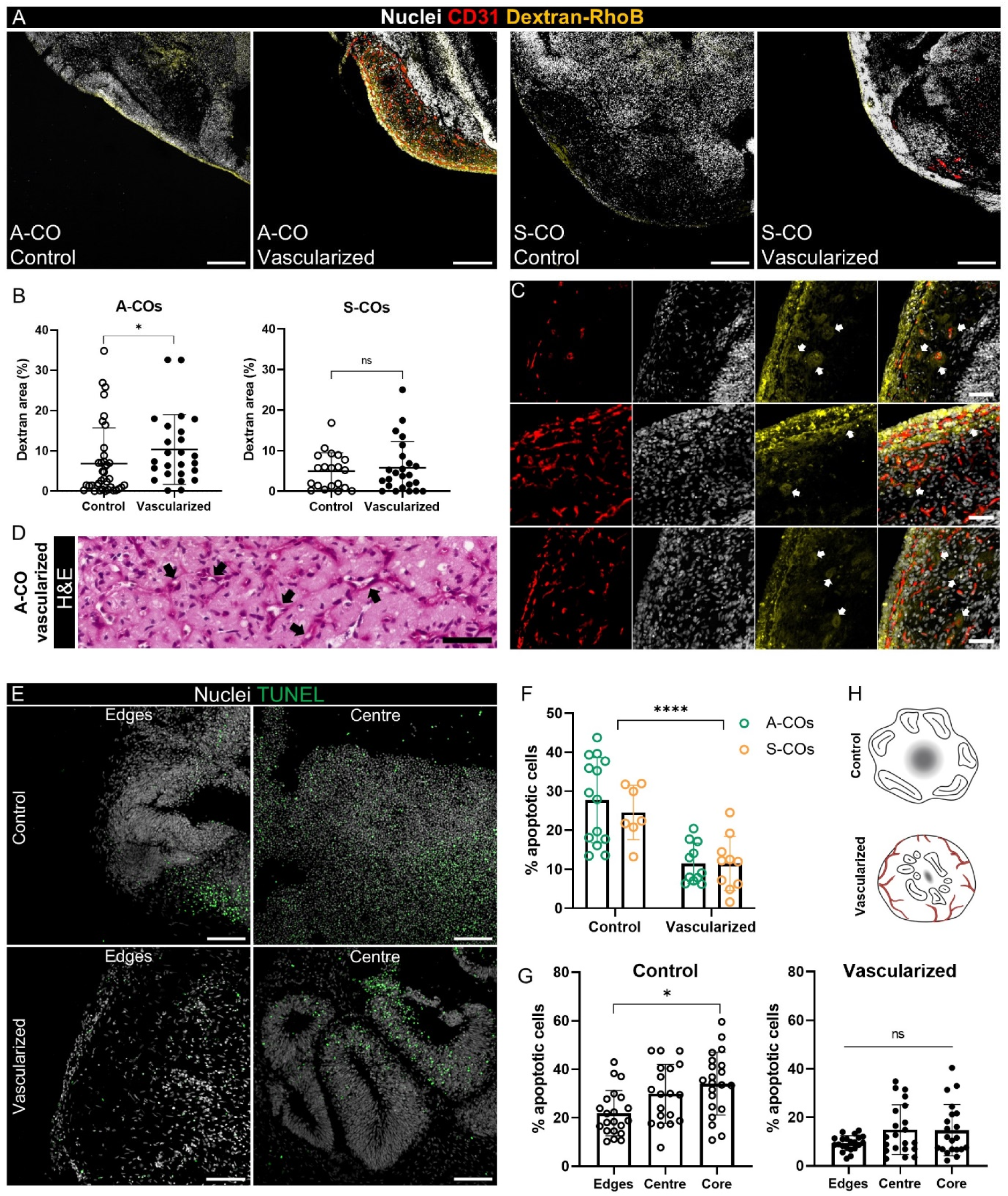
Vascularization of cerebral organoids improves media diffusion through the tissue and reduces the necrotic core size. (A) Confocal microscopy images showing 70 KDa dextran-Rhodamine B (d-RhoB) penetration (yellow) into control and vascularized A– and S-COs. The sections were co-stained with CD31 (red) and nuclei (white). Similar areas across the organoids are shown for consistency. Scale bars: 200 µm. (B) Percentage of d-RhoB penetration (%) normalized by the area of the organoid region analyzed in control and vascularized COs (A-COs on the left and S-COs on the right). Mean ± SD, N = 20-38 (areas analyzed); four organoids per batch, two batches per cell line. Mann-Whitney test; ns = not significant (p > 0.05), *p < 0.05. (C) Panel of confocal images showing representative areas of vascularized A-COs sections in which the d-RhoB solution penetrated deeper into the higher endothelial areas (yellow), leaving halo-shape traces (white arrows) surrounding regions of the endothelial networks (CD31+, red). Scale bars: 50 µm. (D) Paraffin-embedded sections of vascularized A-COs stained with Hematoxylin and Eosin (H&E). Black arrows indicate the presence of lumens within the endothelial networks. Scale bar: 50 µm. (E) Confocal images of representative areas (edge and center) of control and vascularized COs stained for apoptotic cells using the TUNEL assay (green; nuclei in white). Scale bars: 100 µm. (F) Average percentage of apoptotic cells (%) across the central section of control and vascularized COs, for both iPSC lines (A-iPSC, green; S-iPSCs, yellow). Mean ± SD, N = 7-14, four or three batches per iPSC line. Two-way ANOVA; vascularization effect: ****p < 0.0001. (G) Percentage of apoptotic cells (%) across different areas (edges, center, core) of the central section of control and vascularized COs of both A and S iPSC lines. The core is defined as the region of highest cellular density within the section (determined by the higher number of nuclei) typically located in the center but not exclusively. On the left, control COs (white-filled dots) and on the right vascularized COs (black-filled dots). Mean ± SD, N = 20-21, four or three independent batches, two different iPSC lines. Control COs: One-way ANOVA, asterisk showing Tukey’s post hoc comparisons; *p < 0.05. Vascularized COs: Brown-Forsythe and Welch ANOVA test and Games-Howell’s post hoc comparisons; ns = not significant (p > 0.05). (H) Graphical representation of the necrotic core size (grey diffuse circle) between control and vascularized organoids (endothelial networks in red). Neural rosettes (represented as elliptical shapes with a straight line in the middle) are represented towards the organoid edges in control COs, while in the vascularized COs, they are more central, as observed during the TUNEL assay. Created with Biorender.com.

To determine whether the presence of endothelial networks was associated with improved survival, we compared the apoptotic cell levels in control and vascularized COs using a TUNEL assay (Figure 7E). For both iPSC lines, the percentage of apoptotic cells was significantly reduced by almost three-fold when the tissue was vascularized using our adapted CO strategy (Figure 7F). By analyzing the different areas across the sections, we observed a different distribution of apoptotic levels. For instance, in control organoids the increase in cellular death was especially noticeable in the core, which is the more densely populated region typically located at the center, known for this reason as the necrotic core. In the case of vascularized COs, the apoptotic cell levels were more uniformly distributed across the tissue (Figure 7E, H). Neural rosettes exhibited diminished apoptosis levels. In control COs, these structures were generally positioned near the edges, whereas in vascularized COs, they tended to be situated in the central regions (Figure 7E, G).

## DISCUSSION

In this study, we present a simplified vascularization approach for COs to model the human cerebrovasculature *in vitro*. Our strategy generates multicellular endothelial networks both on the surface and within neural tissue and essentially mimics key aspects of the BBB creating a microphysiological system that is suitable for diverse and translational research applications. Compared to controls, our vascularization strategy improved CO tissue viability without disrupting organoid growth or cell differentiation. Moreover, organoids with a higher endothelial network density tend to be more uniform in circularity and size, suggesting that the networks may act as a “vascular mesh” that physically supports the organoids without altering tissue stiffness.

Recent protocol adaptations have grown organoids in the absence of ECM biomaterials ^8,56–59^; however, we sought to leverage the organoid embedding step to incorporate ECs from an early developmental stage so that they synergistically develop together for 30+ days. Previous studies have encapsulated COs at different developmental stages with iPSC-derived ECs^38,39^. Here, we selected HBMVECs given their phenotypic relevance and tissue specificity^50^. Neural tissue expansion, in contrast to matrix contraction, facilitates HBMVEC integration, mimicking the *in vivo* process of central nervous system neovascularization, in which endothelial sprouts growing from mesodermal angioblasts penetrate the developing neural tube^60^. The versatility of this strategy may also be used to incorporate other cell types that are naturally absent in COs, such as microglia^61^. In addition, we demonstrated that, beyond structural support, biomaterial properties can be tuned to control the EC invasion rate and network morphology. We found that EC network formation improved with lower concentrations of hydrogel; however, Geltrex^TM^ reached a critical setting point at 60% and could not be diluted further without compromising the stability of the embedding droplet during dynamic cultures. Further studies with more viscoelastic and tunable biomaterials with defined compositions (i.e., fibrin, gelatin methacrylate, or hyaluronic acid^62–64^) would be interesting to enhance cellular motility and minimize the batch-to-batch variability associated with basement membrane-derived biomaterials, like Matrigel or Geltrex^TM^ ^65^.

In unguided cerebral organoid protocols, the spontaneous differentiation of (peri)vascular cells has been reported without any mesodermal/angiogenic stimulation^66,67^. To control this and to offer reproducible induction, we added ECG media and VEGF during the expansion and maturation of COs. ECG media contain basal fibroblast growth factor (bFGF) which promotes mesoderm induction, especially in the presence of BMP-4, which might be secreted from developing cerebral tissue ^68,69^. Simultaneously, maturation media^9^ include vitamin A, a precursor of retinoic acid, which plays an important role in vasculogenesis^70,71^. Mesodermal progenitors were confirmed to be absent before the expansion stage, implying that endogenous endothelial differentiation occurs after this step following addition of ECG media and VEGF. Differentiation and proliferation of ECs are promoted by a positive feedback loop from VEGF^72^ that promotes the production of other pro-angiogenic proteins, such as MMP-9. The overexpression of these proteinases likely explains the incremental kinetics of Geltrex^TM^ degradation in encapsulated HBMVECs stimulated with VEGF and the initial faster growth of BECs MM+VEGF organoids. We also showed differences between two commercially available iPSC lines, one with a higher (A-iPSCs) and one with a lower (S-iPSCs) ECs differentiation rate. However, inhibiting the SMAD pathway to generate CortiCOs reduced endogenous EC differentiation in A-organoids and prevented mesodermal induction by mixed media or VEGF. We also demonstrated that the vascularization strategy was successful with iPSC-derived ECs, which opens the opportunity to generate isogenic, patient-derived vascularized COs. However, a critical consideration for the wider uptake of this method is the innate variability between iPSC lines, as reported in the literature^73,74^.

Perfusion of 3D organoid systems remains at the cutting edge of model development, employing microfluidic systems to generate a viable 3D network for the entire microtissue^49^. In this study, we attempted to validate the lumenization and potential perfusability of this vascularized tissue. Our study suggests that there is an open, leaky network of ECs, facilitating the internalization of dextran into the tissue, and media diffusion to ameliorate apoptotic tissue regions. Although perfusion of the tissue is desirable, the presence of ECs throughout the tissue may also work to reduce apoptosis via paracrine signaling^37,66,75–77^; however, the development of more advanced systems is required to further assess and stimulate organoid perfusability.

Overall, we developed a vascularized cerebral organoid model using commercially available cells and reagents that reflect key aspects of the human cerebrovasculature. Beyond cerebral tissue, our approach could be used to vascularize organoids of other tissues as well. This model fulfills a crucial requirement in the field by providing an essential platform for the exploration of cerebrovascular diseases mechanisms, development of therapeutics interventions, and modelling of the BBB. This model is expected to have a significant impact on the study of cerebral organoids and aid in the advancement of patient-specific MPS, thereby enhancing personalized medicine.

### Limitations of the study

There are limitations to this study that we hope can be addressed with further characterization of the model or by the wider adoption of our methodology within the organoid field. A key limitation in understanding the effect of vascularization on the maturation and organization of neural tissue is the cessation of CO culture at 40 days, as longer cultures would provide further insights into tissue maturation and similarities with human neurodevelopment. Another major limitation was the perfusion assessment, which requires a dynamic system to demonstrate network functionality, together with assays like trans-endothelial electric resistance (TEER). During neovascularization, flow induces endothelial lumenization and the expression of tight junctions. We believe that by coupling our pre-vascularized organoids to a vascularity on a chip, we will stimulate connectivity and further recapitulate functional neuronal tissue. Finally, due to the variability between iPSC lines, the wider adoption of this strategy will require validation studies to maintain reproducibility.

## STAR★Methods

KEY RESOURCES TABLE.

| REAGENT or RESOURCE | SOURCE | IDENTIFIER |
| --- | --- | --- |
| **Antibodies** |  |  |
| Mouse anti-human CD31 (PECAM-1) | Bio-Rad | Cat# MCA1738; RRID: AB_322710 |
| Rabbit anti-βIII-tubulin (βIIITUB) | Sigma-Aldrich | Cat# T3952; RRID: AB_1841226 |
| Mouse anti-βIII-tubulin (βIIITUB) | Sigma-Aldrich | Cat# T5293; RRID: AB_477580 |
| Rabbit anti-SOX2 | Invitrogen / Thermo Fisher Scientific | Cat# PA1-094; RRID: AB_2539862 |
| Mouse anti-PAX6 | Invitrogen / Thermo Fisher Scientific | Cat# MA1-109, RRID: AB_2536820 |
| Rabbit anti-human Collagen IV | Invitrogen / Thermo Fisher Scientific | Cat# PA5-95188, RRID: AB_2806993 |
| Mouse anti-human Laminin α5 | Invitrogen / Thermo Fisher Scientific | Cat# MA5-24649, RRID: AB_2637264 |
| Rabbit anti-Glial Fibrillary Acidic Protein (GFAP) | Agilent Dako | Cat# Z033401-2 |
| Goat anti-human brachyury (963427) | Bio-technie / R&D Systems | Included in the Human Pluripotent Stem Cell Functional Identification Kit; Cat# SC027B |
| Goat anti-human Otx2 (963273) |  |  |
| Goat anti-human SOX17 (963121) |  |  |
| Rabbit anti-VE-Cadherin | Cell Signaling Technology | Cat# 2500, RRID: AB_10839118 |
| Rabbit anti-PDGFRβ | Invitrogen / Thermo Fisher Scientific | Cat# MA5-15143, RRID: AB_10985851 |
| Mouse anti-α Smooth Muscle Actin (αSMA) | Abcam | Cat# ab7817, RRID: AB_262054 |
| Mouse anti-Human SSEA4 (A24867) | Thermo Fisher Scientific | Included in the Pluripotent Stem Cell Immunocytochemistry Kit (OCT4, SSEA4); Cat# A25526 |
| Rabbit anti-Human OCT4 (A24866) |  |  |
| Anti-human CD31 PerCP/Cyanine5.5 | BioLegend | Cat# 303132 |
| Donkey anti-mouse IgG (H+L) (Alexa Fluor™ 488) | Invitrogen / Thermo Fisher Scientific | Cat# A-21202, RRID: AB_141607 |
| Donkey anti-Rabbit IgG (H+L) (Alexa Fluor™ 594) | Invitrogen / Thermo Fisher Scientific | Cat# A-21207, RRID: AB_141637 |
| Goat anti-rabbit IgG (H+L) (Alexa Fluor™ 647) | Invitrogen / Thermo Fisher Scientific | Cat# A32733, RRID: AB_2633282 |
| Goat anti-mouse IgG (H+L) (Alexa Fluor™ 647) | Invitrogen / Thermo Fisher Scientific | Cat# A-21236, RRID: AB_2535805 |
| Donkey Anti-Goat IgG H&L (Alexa Fluor™ 647) | Abcam | Cat# ab150131; RRID: AB_2732857 |
| **Chemicals, peptides, and recombinant proteins** |  |  |
| Essential 8™ Flex Medium Kit | Thermo Fisher Scientific | Cat# A2858501 |
| Geltrex™ Reduced-Growth Factor Basement-Membrane Matrix, LDEV-free, stem-cell qualified | Thermo Fisher Scientific | Cat# A1413302 |
| RevitaCell™ Supplement (100X) | Gibco / Thermo Fisher Scientific | Cat# A2644501 |
| Gentle Cell Dissociation Reagent | STEMCELL technologies | Cat# 100-0485 |
| KnockOut™ DMEM | Gibco / Thermo Fisher Scientific | Cat# 10829018 |
| Endothelial Cell Growth Medium | PromoCell | Cat# C-22110 |
| Gelatin, from porcine skin | Sigma-Aldrich | Cat# G2500-500G |
| Trypsin-EDTA Solution | Sigma-Aldrich | Cat# T4049 |
| Dimethyl sulfoxide (DMSO) | Sigma-Aldrich | Cat# D2650-100ML |
| Human Recombinant VEGF-165 | STEMCELL technologies | Cat# 78073 |
| Calcein AM | Invitrogen / Thermo Fisher Scientific | Cat# C3100MP |
| OCT compound | VWR Chemicals | Cat# 361603E |
| Fluoromount™ Aqueous Mounting Medium | Sigma-Aldrich | Cat# F4680-25ML |
| D-Sucrose | Thermo Fisher Scientific | Cat# BP220-1 |
| Hanks' Balanced Saline Solution | Sigma-Aldrich | Cat# H9269-500ML |
| Dulbecco's Phosphate Buffered Saline | Sigma-Aldrich | Cat# D8537-500ML |
| DMEM/F-12, HEPES | Gibco / Thermo Fisher Scientific | Cat# 11574546 |
| KnockOut™ Serum replacement | Gibco / Thermo Fisher Scientific | Cat# 10829220 |
| Fetal Bovine Serum | Sigma-Aldrich | Cat# F7524 |
| GlutaMAX™ Supplement | Gibco / Thermo Fisher Scientific | Cat# 35050061 |
| MEM Non-Essential Amino Acids Solution (100X) | Gibco / Thermo Fisher Scientific | Cat# 11140050 |
| Penicillin-Streptomycin | Sigma-Aldrich | Cat# P0781 |
| 2-Mercaptoethanol | Sigma-Aldrich | Cat# M3148 |
| Recombinant Human FGF2 (145 aa) Protein | R&D Systems | Cat# 3718-FB |
| Y-27632 (Dihydrochloride) | STEMCELL Technologies | Cat# 72302 |
| B-27™ Supplement (50X), minus vitamin A | Gibco / Thermo Fisher Scientific | Cat# 12587010 |
| Heparin sodium salt from porcine intestinal mucosa | Sigma-Aldrich | Cat# H3149-10KU |
| SB 431542 hydrate | Sigma-Aldrich | Cat# S4317-5MG |
| LDN193189 hydrochloride | Sigma-Aldrich | Cat# SML0559-5MG |
| Advanced DMEM/F12 | Gibco / Thermo Fisher Scientific | Cat# 11540446 |
| N-2 supplement | Thermo Fisher Scientific | Cat# 11520536 |
| Neurobasal™ Medium | Gibco / Thermo Fisher Scientific | Cat# 11570556 |
| B27 Supplement 50X serum free | Thermo Fisher Scientific | Cat# 11530536 |
| Insulin solution human | Sigma-Aldrich | Cat# I9278-5ML |
| Recombinant human NT3 | PeproTech / Thermo Fisher Scientific | Cat# 450-03 |
| Human/Mouse/Rat BDNF Recombinant Protein | PeproTech / Thermo Fisher Scientific | Cat# 450-02 |
| Anti-Adherence Rinsing Solution | STEMCELL Technologies | Cat# 07010 |
| Advanced DMEM | Thermo Fisher Scientific | Cat# 12491015 |
| L-Ascorbic Acid | Sigma-Aldrich | Cat# A5960-25G |
| Laduviglusib (CHIR-99021) | MedChemExpress / Thermo Fisher Scientific | Cat# 16041282 |
| Invitrogen™ Dynabeads™ Sheep Anti-Mouse IgG | Invitrogen / Thermo Fisher Scientific | Cat# 10630514 |
| Rhodamine B isothiocyanate–Dextran | Sigma-Aldrich | Cat# R9379-250MG |
| Agarose, low gelling temperature | Sigma-Aldrich | Cat# A9414-10G |
| Hoechst 33342 solution, 20 mM | Thermo Fisher Scientific | Cat# 62249 |
| 7-Aminoactinomycin D (7-AAD) | Thermo Fisher Scientific | Cat# A1310 |
| Paraformaldehyde | Sigma-Aldrich | Cat# 158127-500G |
| Ethanol | Honeywell | Cat# 32221 |
| Bovine serum albumin | Sigma-Aldrich | Cat# A2153-50G |
| Triton™ X-100 | Sigma-Aldrich | Cat# T878-250ML |
| Trisodium citrate dihydrate | Sigma-Aldrich | Cat# S1804-500G |
| Tween™ 20 | MP biomedical | Cat# TWEEN201 |
| NaCl | Sigma-Aldrich | Cat# S9888 |
| MgCl<sub>2</sub> | Sigma-Aldrich | Cat# M8266 |
| HEPES | Sigma-Aldrich | Cat# H4034 |
| Xylenes 98% | Thermo Fisher Scientific | Cat# 383930050 |
| DPX mountant for histology | Sigma-Aldrich | Cat# 06522-500ML |
| Hematoxylin solution (Mayer's hemalum solution) | Sigma-Aldrich | Cat# 1.09249-500ML |
| Eosin Y solution | Thermo Fisher Scientific | Cat# 10562614 |
| Tetrahydrofuran | Sigma-Aldrich | Cat# 360589-500ML |
| Triethylamine | Sigma-Aldrich | Cat# 471283 |
| Dibenzyl ether | Thermo Fisher Scientific | Cat# 10113630 |
| **Critical commercial assays** |  |  |
| STEMdiff™ Cerebral Organoid Kit | STEMCELL Technologies | Cat# 08570 |
| Pluripotent Stem Cell Immunocytochemistry Kit (OCT4, SSEA4) | Thermo Fisher Scientific | Cat# A25526 |
| Included in the Human Pluripotent Stem Cell Functional Identification Kit | Bio-technie / R&D Systems | Cat# SC027B |
| Human MMP-9 ELISA Kit | Proteintech | Cat# KE00164 |
| Human VEGF ELISA Kit - Quantikine | Bio-technie / R&D Systems | Cat# SVE00 |
| Trichrome Stain (Masson) Kit | Sigma-Aldrich | Cat# HT15 |
| Click-iT™ Plus TUNEL Assay Kits for In Situ Apoptosis Detection | Invitrogen / Thermo Fisher Scientific | Cat# 15350244 |
| MycoStrip® Mycoplasma Detection Kit | InvivoGen | Cat# rep-mysnc-50 |

| **Experimental models: Cell lines** |  |  |
| --- | --- | --- |
| Gibco™ Human Episomal iPSC Line (A-iPSCs) | Gibco / Thermo Fisher Scientific | Cat# A18945 |
| Healthy Control Human iPSC Line SCTi003-A (S-iPSCs) | STEMCELL technologies | Cat# 200-0511 |
| Human brain microvascular endothelial cells | Innoprot | Cat# P10361 |
| **Software and algorithms** |  |  |
| GraphPad Prism 8.0.2 | GraphPad | https://www.graphpad.com/scientificsoftware/prism/ |
| Angiotool | Public (NIH) | https://ccrod.cancer.gov/confluence/display/ROB2/Downloads |
| ImageJ 1.54f | Public (NIH) | https://imagej.net/ij/ |
| Optics11 DataViewer V2.5.0 | Optics11 life | https://www.optics11life.com/products/pavone-nanoindenter/#applications |
| QuPath-0.4.3 | Public (GNU) | https://qupath.github.io/ |
| Aperio ImageScope | Leica Biosystems | https://www.leicabiosystems.com/digital-pathology/manage/aperio-imagescope/ |
| Fluoview software FV31S | Evident | https://evidentscientific.com/en/software/cellsens |
| Fusion software | OXFORD instruments | https://fusion-benchtop-software-guide.scrollhelp.site/fusionum/ |
| FlowJo v10 | BD Biosciences | https://www.flowjo.com/flowjo/download |

### Solutions and media compositions

| Name | Composition |
| --- | --- |
| Permeabilization buffer | 300 µM sucrose, 50 µM NaCl, 6.3 µM MgCl<sub>2</sub>, 20 µM HEPES, 0.5% Triton X-100, pH 7.2 in distilled water |
| Phase I media | DMEM F-12 with HEPES, 20% KOSR, 3% FBS, 1% Glutamax, 1% MEM-NEAA, 1% Pen/Strep, 0.1 mM 2-mercaptoethanol, 4 ng/mL fibroblast growth factor 2 (FGF2) and 50 µM ROCK inhibitor |
| Neural induction media | DMEM F-12 with HEPES, 20% KOSR, 1% Glutamax, 1% MEM-NEAA, 1% Pen/Strep, 0.1 mM 2-mercaptoethanol, 10 µM SB431542 and 100 nM LDN-193189 |
| EB1 medium | Advanced DMEM/F12, 1% N2, 1% Glutamax, 1% Pen/Strep, 0.5% B27 without vitamin A, 4 µg/mL Heparin |
| EB2 media | 50% Advanced DMEM/F12, 50% Neurobasal media, 1% N2, 1% Pen/Strep, 1% MEM-NEAA, 0.5% Glutamax, 0.5% B27 without vitamin A, 2.5 µg/mL insulin) |
| EB3 media | 50% Advanced DMEM/F12, 50% Neurobasal, 0.5% N2, 1% Pen/Strep, 0.5% MEM-NEAA, 1% Glutamax, 1% B27, 2-mercaptoethanol, 5 µg/mL insulin, 20 ng/mL NT3 and 20 ng/mL BDNF |
| LARS media | Advanced DMEM, 60 µg/mL ascorbic acid, 2.5 mM Glutamax |
| Sodium citrate buffer | 10 mM sodium citrate, 0.05% Tween 20, pH 6, in PBS |
| FACS buffer | 2% FBS in PBS |

### Resource availability

### Lead contact

Further information and requests for resources and reagents should be directed to and will be fulfilled by the lead contact, Mihai Lomora.

### Materials availability

The Gibco™ Human Episomal iPSC Line (hPSCreg name <u>TMOi001-A</u>, A18945, Female) has been discontinued but is available in our biobank and maintained by our group. All other materials used in this study are commercially available with no restrictions on availability.

### Data and code availability

This paper analyzes existing, publicly available data. Macros for image analysis can be made available from the lead contact upon request. Any additional information required to reanalyze the data reported in this study is available from the lead contact upon request.

### Experimental model and study participant details

#### Cell lines

A-iPSCs (Gibco™ Human Episomal iPSC Line, Cat# A18945, Female) and S-iPSCs (STEMCELL TECHNOLOGIES Healthy Control Human iPSC Line SCTi003-A, Cat# 200-0511, Female) were purchased commercially, while Zombie Green iPSCs were gifted from Dr Meenakshi Suku, Prof Michael Monaghan (Trinity College Dublin) and Prof Lesley Forester (University of Edinburgh) and are originally described here^78^. Primary HBMVECs, from a pool of multiple donors were purchased from Innoprot (Cat# P10361).

## EXPERIMENTAL MODEL DETAILS

### General cell culture conditions

All cells were maintained at 5% CO_2_ and 37 °C and handled under standard biosafety level 2 (BSL-2) sterile conditions. Routine mycoplasma inspections were conducted to ensure cell integrity following the manufacturer’s protocol for the MycoStrip® Mycoplasma Detection Kit (InvivoGen).

### iPSC culture and cyropreservation

All iPSC lines were cultured on Geltrex^TM^-coated plates (1:100 dilution in DMEM KnockOut^TM^ media, incubated for 2h at 37 °C) and maintained using Essential 8^TM^ Flex Medium (E8 medium). iPSCs were revived from liquid nitrogen by gently adding the contents of the cryovial into 9 mL of DMEM KnockOut^TM^ media, centrifuging at 200 × g for 3 min, and resuspending the pellet carefully to preserve the cell aggregates in E8 media supplemented with 1X RevitaCell^TM^ Supplement for the first 24h only. The medium was exchanged daily, and the colonies were monitored for signs of differentiation. Differentiated cell areas were identified under a CytoSMART^TM^ Lux 2 (AXION Biosystems) microscope inside a biosafety cabinet and manually removed using a pipette tip. At 60% confluency, and before colonies started merging, iPSCs were passaged by detaching the colonies with Gentle Cell Dissociating Reagent (GCDR) for 8 min at 37 °C. The iPSCs were then gently lifted with fresh E8 to maintain cell aggregates, before being transferred to a new plate pre-coated with Geltrex^TM^ and fresh media. Split ratios varied depending on cell confluency, ranging from 1:4 to 1:10. For cryopreservation, iPSCs were incubated for 1h before cell detachment in fresh media supplemented with 1X Revitacell^TM^. Cells were then manually detached using a cell scraper, collected in the same medium, and pelleted for 3 min at 200 × g. The cell pellet was then resuspended in 90% knockout serum replacement medium (KOSR) and 10% dimethyl sulfoxide (DMSO) solution for long-term storage in liquid nitrogen. iPSCs were always cultured for at least three passages before the start of differentiation, and all organoids were generated from passages 9-11.

### Endothelial cells culture

Endothelial cells (ECs) were cultured in Endothelial Cell Growth Medium (ECG medium) on cultureware coated with 1% gelatin. To prepare the coating, gelatin was dissolved in distilled water, autoclaved and filtered through a 0.02 µm syringe filter. The 1% gelatin solution was added to fully cover the cultureware for 1h at RT and then removed to allow it to dry for 1h more at RT. ECs were thawed using pre-warmed ECG media, seeded, and maintained with a full media exchange every 2-3 days. Cultures at 80-90% confluence were propagated to a split ratio 1:3-1:4. For passaging, cells were washed with phosphate buffer saline (PBS) and trypsinized with sufficient pre-warmed Trypsin-EDTA 0.25% to cover the culture surface for approximately 1 min. Cells were detached and collected in ECG media and centrifuged at 500 × g during 5 min at RT. The resulting cell pellet was resuspended in ECG media at the desired cell density for passage and seeded onto fresh gelatin-coated cultureware. For cryopreservation, the cell pellet was resuspended in a cell freezing medium composed of 70% ECG medium, 20% fetal bovine serum (FBS) and 10% DMSO at a density of 2 million cells/mL. The cells were stored in a liquid nitrogen tank for long-term storage. For HBMVECs, the first biobank was generated at passage 5 for future propagation and a second biobank was generated at passage 7 for experimental use. HBMVECs were seeded and cultured for one passage before their use, ensuring that all experiments conducted in this study were performed at passage 8.

## METHOD DETAILS

### HBMVECs network formation

HBMVECs were cultured and trypsinized as previously described, then diluted to the appropriate cell density to achieve 100,000 cells per well in a 96-well plate. Different Geltrex^TM^ concentrations (80%, 60%, 50%, and 40%) were prepared by adjusting the Geltrex^TM^: Media + Cells solution ratio while keeping the cell density constant. To minimize batch-to-batch variability, three Geltrex^TM^ vials were pooled and the specific pool used for each experiment was recorded. Geltrex^TM^ was thawed on ice, and the cell solution was kept on ice shortly before mixing to prevent premature hydrogel polymerization. Geltrex^TM^-encapsulated cells were seeded in a 96-well assay plate (Costar/Corning, Cat# 3603) at a final volume of 50 µL, then polymerized at 37 °C for 30 min (80%, 60% Geltrex^TM^) or 45 min (50%, 40%). After the polymerization, 200 µL of ECG media was added on top of the hydrogels and replaced every other day. When required, the ECG medium was supplemented with 50 ng/mL of vascular endothelial growth factor 165 (VEGF) to promote angiogenesis. To determine the optimal organoid and HBMVECs co-culture media, ECG media was mixed with the maturation media of the STEMdiff™ Cerebral Organoid Kit (STEMCELL technologies, Cat# 08570) at 1:0, 1:1, 1:3, 1:7, and 0:1 (respectively) always supplemented with 50 ng/mL of VEGF. The media was exchanged every other day. To determine the optimal VEGF dosage, ECG media was supplemented with 25 or 50 ng/mL of VEGF, and the media was exchanged every 2 or 4 days. On day 10, the vascular networks were stained with Calcein AM (1:1000 dilution) in Hanks’ Balanced Saline Solution at 37 °C under agitation on an orbital shaker (85 rpm) for 30 min. High-throughput imaging was performed using the Operetta CLS system (Revvity), and the images were analyzed using AngioTool64 v0.6a to determine parameters such as the total and average vessel length, junction density, total number of end points, and lacunarity^79^.

### iPSC quality control assessment

The pluripotency of iPSC lines was confirmed by immunostaining for the pluripotency markers SEEA4 and OCT4 using antibodies included in the Pluripotent Stem Cell Immunocytochemistry Kit (Thermo Fisher Scientific, Cat# A25526). iPSC colonies were grown on glass coverslips until reaching 50-70% confluence and fixed with 4% paraformaldehyde (PFA) for immunostaining. Simultaneously, pluripotency was assessed using the Human Pluripotent Stem Cell Functional Identification Kit (R&D Systems, SC027B) according to manufacturer’s instructions. In short, colonies were grown on glass coverslips for 2 days, as more efficient differentiation was obtained at a lower confluency than the recommended (∼30%). Differentiation into ectodermal, mesodermal or endodermal progenitors was induced. Differentiated cells were fixed with 4% PFA and stained for the specific germinal layer markers OTX2, brachyury and SOX17 (ectoderm, mesoderm and endoderm, respectively). For both pluripotent assays, cells were permeabilized and blocked with a 0.3% Triton X-100 and 1% BSA in PBS solution for 1h at RT. Primary antibodies were diluted in the same solution (OTX2, brachyury, SOX17 at 1:50; SSEA4, OCT4 at 1:200) and the wells were incubated for 3h at RT. Cells were washed with 1% BSA in PBS and incubated with the corresponding secondary antibody diluted 1:500 in the same solution for 1h at RT in the dark. Cells were washed again with 1% BSA, and the nuclei were stained with 1:2000 Hoechst in PBS for 5 min at RT. Wells were washed with PBS and coverslips were mounted onto glass microscopy slides with Fluoromount™ Aqueous Mounting Medium for further confocal microscopy imaging.

### Generation of cerebral organoids

Cerebral organoids (COs) were generated based on the protocol described by Lancaster et al^9^, adapted by the STEMdiff™ Cerebral Organoid Kit. Briefly, iPSCs were visually inspected to ensure optimal colony morphology (compact, sharp edges) and <1 % differentiated cells. On day 0, 9,000 iPSCs were plated in low-adherence 96-well round-bottom plates (Sarstedt, Cat# 83.3925) in Embryoid Body (EB) formation medium supplemented with 10 μM Y-27632 (Rho-associated coiled-coil containing protein kinase (ROCK) inhibitor). The plates were left undisturbed for the first 24h, with additional EB formation media (without the ROCK inhibitor) added on days 2 and day 4. At this stage, the EBs typically reached a diameter of >300 µm. On day 5, individual EBs were transferred to a 24-well plate containing induction media using a cut 200 μL micropipette tip. To minimize medium carryover, the organoids were allowed to sink to the end of the tip before transfer. EBs were incubated in induction media until they exhibited smooth and optically translucent edges (neuroepithelium development) typically between days 7-9 (iPSC line dependent). For consistency, all organoids were maintained in the induction stage until day 8. On the encapsulation day, embedding sheets were prepared by pressing the bottom of a 2 mL Eppendorf tube onto parafilm pieces placed onto a micropipette tip holder to create small concavities that could hold a small amount of liquid. The sheets were sterilized by immersion in 100% ethanol for 15 min, followed by UV exposure for 30 minutes, and then allowed to dry inside the biosafety cabinet. Each concavity of the sterilized sheet could accommodate one CO embedded in a 25 µl Geltrex^TM^ droplet and polymerized at 37 °C for 30 min. The droplets were collected in a 6-well plate by lifting the embedding sheets with sterile tweezers and gently sliding them down with PBS. Once collected, PBS was replaced with expansion media, and the organoids were incubated for 3 days. After this period, the media were switched to maturation media and transitioned to a dynamic culture by placing the plate on a CO_2_-resistant orbital shaker (Thermo Fisher Scientific, Cat# 88881104B) set at 85 rpm in a 37 °C incubator. The media were fully replaced every 3-4 days until day 40, when organoids were collected for downstream analysis. All cultureware used for COs culture were treated with anti-adherence rinsing solution (AARS) and washed with PBS before use.

### Generation of cortical organoids

Cortical organoids (CorticOs) were generated following an in-house protocol developed by Chan et al. (unpublished). This in-house protocol follows the same protocol as Lancaster et al^9^ but incorporating dual SMAD inhibition during the induction stage^80^. The composition of each medium is detailed in the **Key Resources Table**. As described above, 9,000 iPSCs were aggregated in an AARS-treated low-adherence 96-well plate in Phase I media. Phase I media was added on day 2 and on day 4, with the latter excluding FGF2 and ROCK inhibitor. On day 6, EBs were individually transferred to AARS-treated 24-well plates containing neural induction media as previously described. On day 8, CortiCOs were encapsulated in Geltrex^TM^ as per the COs protocol and transferred to AARS-treated 6 well plates with EB1 medium on an orbital shaker (85 rpm). The media was exchanged every 3 or 4 days, transitioning to EB2 media from day 16 and EB3 media from day 30. Both brain organoid protocols are compared in **Figure S2** of the **Supporting Information**.

### Optimization of the HBMVECs incorporation within cerebral organoids

HBMVECs were incorporated in the organoids culture during the Geltrex^TM^ embedding step (Day 8). In the encapsulation approach, HBMVECs were collected as previously described, brought to the appropriate cell density in cold ECG media, and mixed with Geltrex^TM^ (20% cell solution, 80% Geltrex^TM^) on ice. Organoids were embedded within 25 μL of the mixture, polymerized and collected as previously described. In the surface attachment approach, organoids were embedded in 100% Geltrex^TM^ droplets as previously described. Organoids were individually collected in an AARS-treated 48-well plate, to which the appropriate number of HBMVECs was added. In the surface attachment approach, 2M and 500K cells per organoid were tested. The encapsulation approach was further investigated with 2M, 500k and 50K cells per organoid. Once the HBMVECs were introduced, the expansion and maturation media from the STEMdiff™ Cerebral Organoid Kit were supplemented with 50 ng/mL VEGF. Control organoids were grown (100% Geltrex^TM^, without HBMVECS) in standard media with and without VEGF supplementation. The results of this section are shown in **Figure S1.**

### Encapsulation of HBMVECs into the organoid at different Geltrex^TM^ concentrations

Except as explicitly stated, 50,000 HBMVECs were encapsulated per organoid (+BECs). On day 5 of the organoid protocol, HBMVECS were seeded and cultured as previously described. On the embedding day (day 8), HBMVECs were concentrated to 10 million cells/mL in ECG media and kept cold to prevent premature Geltrex^TM^ polymerization. To obtain a 80% concentration, one part of the cell solution was mixed with four parts of cold Geltrex^TM^. Organoid embedding with 25 µL of the cell-encapsulating hydrogel was performed as previously described. When the organoids were embedded without cells, cold PBS was used to dilute Geltrex^TM^ to maintain the hydrogel concentration. The different Geltrex^TM^ percentages tested (60% and 95%) were obtained by adjusting the ratio between the hydrogel and cell suspension/PBS, while keeping the final mixture at 2 million cells/mL in all conditions. From this point onward, both COs and CortiCOs were cultured in three media conditions: *i)* control media (CTRL, standard organoid-specific media), *ii)* media supplemented with 50 ng/mL VEGF (+VEGF), and *iii)* mixed media (one part ECG media + seven parts organoid media) supplemented with 50 ng/mL VEGF (MM+VEGF).

### Generation of fluorescent iPSC-derived endothelial cells

Fluorescent iPSC-derived endothelial cells (Z-ECs) were differentiated from the Zombie Green iPSC line (Z-iPSCs) using the protocol described by Browne et al.^81^. Z-iPSCs were cultured as previously described until they reached approximately 40% confluency (typically two-three days after passage), when the media was exchanged for LARS media supplemented with 5 μM CHIR99021. On ay 2 and day 4 of differentiation, the media were fully replaced with LARS media supplemented with 50 ng/mL VEGF. Magnetic sorting of the platelet endothelial cell adhesion molecule 1 (PECAM-1 or CD31)-positive cells was performed on day 6 using the EasySep™ Magnet (STEMCELL Technologies, Cat# 18000). Cells were trypsinized with trypsin diluted in PBS (1:2), collected by adding 2% FBS in PBS, and pelleted at 200 × g for 3 min. The cell pellet was resuspended in 2% FBS-PBS containing anti-human CD31 antibody (1:50 dilution) and incubated for 1h at RT in a rotating shaker. Cells were pelleted (200 × g for 3 min), washed with 2% FBS-PBS, pelleted again and then incubated with 7 μL of Dynabeads™ Sheep-Anti Mouse IgG magnetic beads in 2% FBS-PBS for 1h at RT on a rotating shaker. After incubation, CD31-cells were discarded, and CD31+ cells were washed and eluted in ECG media using the magnet stand and a 3 mL syringe (Terumo, Cat# MDSS03SE) acting as a separation column. CD31+ cells were seeded in a cell culture flask coated with 1% gelatin and ECG media supplemented with 5 μM ROCK inhibitor. Z-ECs were cultured and propagated similar to HBMVECs, but with the addition of ROCK inhibitor the day after passaging to improve survival, prior to their use at passage 3 to 4.

### Nanoindentation

Nanoindentation of individual organoids was performed using a PAVONE Nanoindenter (Optics11 Life) equipped with a spherical probe (9.5 μm tip radius, 0.47 N/m stiffness) and a 2.82 geo factor in air (Optics11 Life, serial number PV220197). On day 40, organoids were collected, washed with PBS and immobilized to the bottom of a 6-well plate using a thin layer of 3% low-melting-point agarose in PBS, which was previously dissolved and maintained at 37 °C. Importantly, the organoids were not completely embedded in the agarose layer. PBS was then carefully added to the wells for nanoindentation analysis. Measurements were conducted at RT with a loading/unloading rate of 200 µm/s, a displacement of 60 µm and a dwell time of 2 s at the peak load. Reference surface calibration was performed against the bottom of the well plate. Three measurements were performed for each organoid, at different ROIs. To determine the Young’s modulus, a Hertzian contact model was applied using the Optics11 DataViewer V2.5.0 software (Optics11 Life) with P_max_(%) = 100 and P_max_(%) for the contact point = 20.

### Enzyme-linked immunosorbent assays (ELISAs)

ELISAs were performed on the organoid culture supernatants according to the manufacturer’s protocols for the Human MMP-9 ELISA Kit (Proteintech, Cat# KE00164) and Quantikine Human VEGF ELISA Kit (Bio-techne, Cat# SVE00). Organoid culture supernatants were collected after each media exchange and mixed with FBS (1:20) to stabilize small peptides and cytokines. For VEGF quantification, the supernatants were diluted 200 times for the +BECs MM+VEGF condition and 20 times for the control COs. For MMP-9 quantification, +BECs MM+VEGF supernatants were diluted 10 times, while control organoid supernatants were analyzed undiluted. These dilutions were established after a preliminary test to ensure that the absorbance values remained within the range of the standard curves. Absorbance at 450 nm was measured using a Varioskan® Flash microplate reader (Thermo Fisher Scientific, Cat# N06354). For VEGF quantification, the 570 nm values were measured and subtracted from the 450 nm readings, while for MMP-9, the 630 nm values were subtracted.

### Whole organoid immunostaining

After 40 days of culture, the organoids were washed three times with PBS and fixed with a 4% PFA solution overnight (O/N) at 4 °C. Following fixation, organoids were washed with PBS and incubated for 2h at RT with a permeabilization/blocking solution (5% BSA and 0.5% Triton X-100 in PBS) on a plate shaker. Primary antibodies were diluted at the specified dilutions (1:200 for CD31; 1:500 for βIII-tubulin) in a solution of 1% BSA and 0.1% Triton X-100 in PBS and incubated O/N at 4 °C under agitation. The next day, the organoids were washed three times (fast wash, 5 min, and 10 min under agitation) with a 0.05% Tween 20 in PBS (PBS-T) solution and incubated with the secondary antibody diluted 1:500 in 1% BSA and 0.1% Triton X-100 in PBS solution for 1h at RT under agitation. Finally, the organoids were washed with PBS (fast spin, 5 min, and 10 min) and kept at 4 °C in a glass-bottom 3.5 cm µ-dish (IBIDI, Cat# 81158) prior to confocal imaging. Day 5 and day 8 embryonic bodies were stained following the same procedure but using brachyury and OCT4 primary antibodies (diluted 1:50) and secondary antibodies (diluted 1:250). A complete list of the antibodies used in this study is available in the **Key Resources Table.**

### FDISCO clearing of whole organoid tissue

The tissue was cleared following an adapted FDISCO protocol^82^. Following whole organoid immunostaining, the organoids were incubated for dehydration (24h, 4 °C) in 50%, 70%, 90%, and 100% sequential aqueous tetrahydrofuran (THF) solutions, with the pH adjusted to 9.0 using triethylamine. The organoids were then transferred to 100% dibenzyl ether (DBE) to achieve a tissue-matching optical refractive index of 1.5605-1.5645, and stored in DBE at 4 °C in glass containers. For imaging, the organoids were transferred to a glass-bottom 3.5 cm µ-dish and imaged using an inverted confocal microscope.

### Organoid cryosectioning and cryopreservation

Organoids were fixed with 4% PFA in PBS O/N at 4 °C under agitation. Then, they were equilibrated with a 15% sucrose in PBS solution O/N at 4 °C under agitation and, the following day, transferred to a 30% sucrose in PBS solution, kept O/N at 4 °C under agitation. Once the organoids sank to the bottom of the well, they were embedded in Optimal Cutting Temperature (OCT) mounting media using disposable 15 x 15 mm base molds (Epredia, Cat# 58950), snap-frozen in liquid nitrogen and stored at –70 °C. A DAKEWE 6250 cryostat (DAKEWE) was used to section the organoids at a thickness of 15 μm, ensuring proper tracking of the collected sections to identify regions across the organoids. Cryosections were attached to SuperFrost™ Plus Adhesion microscopy slides (Epredia, Cat# J1830AMNZ) and kept at –20 °C for immediate use or –70 °C for long-term storage.

### Immunohistochemistry

In preparation for immunohistochemistry (IHC), the slides were warmed to RT for 1-2h and left to dry to ensure proper adherence of the organoid sections. A hydrophobic circle was drawn around the sample using a liquid blocker super PAP pen and allowed to dry. For immunostaining of some intracellular markers (i.e. SOX2, PAX6, collagen IV, and laminin), an additional antigen retrieval step was performed by immersing the slides in sodium citrate buffer and heating them in a food steamer for 20 min. All slides were washed with PBS and permeabilized with cold permeabilization buffer for 5 min at RT, followed by blocking with 1% BSA in PBS for 30 min at RT. Primary antibodies were diluted in the same blocking buffer at the indicated concentrations: αSMA (1:100); CD31, PDGFRβ, SOX2, PAX6, GFAP, Laminin (1:200); VE-CAD (1:400); and βIIITUB, Collagen IV (1:500). The primary antibody solution was added to the slides, covered with parafilm to prevent drying, and incubated at 4 °C O/N. The next day, the slides were washed thrice with PBS-T and 1:500 secondary antibodies, diluted in blocking buffer, followed by incubation for 1h at RT. After secondary antibody incubation, the slides were washed three times with PBS and incubated in a 1:2000 Hoechst in PBS solution for 5 min at RT for nuclei staining. Slides were sealed with a coverslip using Fluoromount™ Aqueous Mounting Medium and dried O/N at RT. The slides were further sealed with nail polish and kept at 4 °C until imaging using a confocal microscope. A complete list of antibodies used in this study is available in the **Key Resources Table.**

### Flow cytometry

iPSC-derived ECs were cultured and propagated as previously described. Cells were detached with 1:2 0.25% trypsin-EDTA in PBS solution and lifted with excess of ECG medium. Subsequent steps were performed on ice or at 4 °C. Cells were divided into 1 million per tube, pelleted by centrifugation (800× g, 5 min), and incubated in FACS buffer containing PerCP-Cy5.5 anti-human CD31 antibody (5 µL per sample) for 1 h at 4 °C on an Eppendorf tube rotator. After incubation, the cells were pelleted, washed twice with PBS, and resuspended in FACS buffer for analysis. The viability dye 7-AAD was added before running the sample to distinguish between live and dead cells. The controls included unstained and single-stained cells for compensation. Flow cytometry was performed using a BD Accuri™ C6 Plus (BD Biosciences) cytometer, and data were analyzed with FlowJo v10.

### Tubular assay

Z-ECs were cultured as previously described. To assess tubular network formation, 60 µl of 100% Geltrex^TM^ was added to each well of a 96-well assay plate (Costar/Corning, Cat# 3603) and incubated at 37 °C for 30 min to allow polymerization. Then, 20,000 cells were seeded onto the matrix in 200 µL of ECG media supplemented with 50 ng/mL VEGF. After 24 h, fluorescent tubular structures were imaged using an Olympus IX81 inverted scanning fluorescence microscope (Evident).

### Paraffin-embedded tissue processing

On day 40, the vascularized organoids were fixed with 4% PFA O/N at 4 °C. The tissue was then embedded in paraffin wax at 58 °C in an Excelsior AS Tissue Processor (Thermo Fisher Scientific). Tissue sections were cut to a thickness of 5 μm using a rotary microtome (Leica RM2235) and mounted on SuperFrost™ Plus Adhesion microscopy slides.

### Histological staining

Cryosection slides were dried at RT for 1-2h and Masson’s trichrome histological staining was performed according to the manufacturer’s protocol (Sigma-Aldrich, Cat# HT15). Paraffin-embedded slides were dewaxed in xylene and rehydrated with water through a sequence of ethanol concentrations (100%, 100%, 90%, and 70%). Slides were submerged for 10 s in hematoxylin solution, and an eosin counterstain was used for 30 s before tissue dehydration in xylene. Slides were mounted with DPX mountant and images were taken using the Ocus®40 slide scanner (Grundium) and processed using QuPath v0.4.3 or ImageScope software (v12.4.6.5003).

### 70 kDa dextran penetration tests

Individual day 40 organoids (control and vascularized) were transferred to a 96-well assay plate (Costar/Corning, Cat# 3603) together with 100 μL of media using a cut tip, and brightfield images were captured to measure the organoid size. The same volume of maturation media containing 70 kDa dextran-Rhodamine B (d-RhoB) was added to each well to achieve a final concentration of 1 mg/mL. Organoids were incubated under the same culture conditions for 10 h (37 °C, orbital shaker 85 rpm). The organoid was then transferred together with 25 μL of the d-RhoB-media solution to another 96-well plate containing 100 μL of PBS. The new plate was shaken for 10 min to ensure the diffusion of the fluorescent solution from the organoid. Following this, the organoid was removed, and the fluorescence of the content of the wells was measured (excitation 570 nm, emission 595 nm) using a Varioskan® Flash microplate reader. Fluorescent values were first normalized to a percentage, considering 100% the fluorescent intensity of 25 μL d-RhoB solution without any organoid. These percentages were then normalized to the area of each organoid. The results of this experiment are shown in **Figure S5.** To visually identify the penetration of 70 kDa dextran across the organoid tissue, day 40 organoids were exposed to the same 1 mg/mL d-RhoB solution in maturation media for three hours at 37 °C under agitation. Finally, the organoids were processed for cryosectioning as previously described. The obtained sections were co-stained for CD31 and nuclei following IHC protocol described above.

### TUNEL assay

The TUNEL assay was performed in sections of the central part of the organoid (determined by counting the total number of sections) following the manufacturer’s protocol of Click-iT™ Plus TUNEL Assay Kit (Invitrogen, Cat# 15350244). Images were acquired using confocal microscopy and analyzed using ImageJ. The total number of nuclei stained by the TUNEL assay was related to the total number of nuclei to determine the percentage of apoptotic cells across all the image fields for each organoid. To determine the apoptotic rate in different regions of the organoids (edges, center and core), image fields were classified based on the part of the organoid they contained, and considering the core as these areas with the highest content of nuclei. The overall percentage of apoptotic cells in these areas was calculated by averaging the individual percentage of each image field.

### Confocal microscopy

Individual organoids (and embryonic bodies) were imaged in a glass-bottom 3.5 cm µ-dish (IBIDI, Cat# 81158). Images were acquired using a FLUOVIEW FV3000 confocal laser scanning microscope (Olympus Scientific Solutions) equipped with 4x, 10x and 20x UPlanSApo objectives (Olympus) and processed using Fluoview version FV31S. Z-stacks were taken every 30-40 μm for day 40 organoids and 7-10 μm for embryonic bodies. Immunostained slides were imaged using a BC43 Andor benchtop confocal microscope (OXFORD instruments), using 10x and 20x objectives, often scanning the entire section using a multilocation montage and a Z-scan of 10-15 μm (images every 2-5 μm). Images were acquired and processed using the Fusion imaging software (OXFORD Instruments).

## QUANTIFICATION AND STATISTICAL ANALYSIS

### Organoid morphology and Geltrex^TM^ degradation rate analysis

During organoid culture, images of growing organoids were acquired during media changes using a phone camera and an Olympus CKX41 inverted phase-contrast microscope (Microscope central, Cat# CKX41-B) equipped with a 4x objective. The scale was corrected using a Neubauer chamber image as a reference image. Images were analyzed using ImageJ v1.54f. Organoid size was quantified by thresholding the brightfield images and using the “Measure particles” function to determine the area and shape indicators (i.e., circularity). For the Geltrex^TM^ degradation rates, the droplets of hydrogel containing HBMVECs were manually contoured, and the area was quantified whenever possible. When comparing the effect of media conditions (CTRL vs. +VEGF vs. MM+VEGF) on +BECs organoids, the area of the organoid was subtracted from the total area of the Geltrex^TM^ droplet to account for the difference in cellular invasion between conditions.

### Fluorescent image analysis

If not explicitly stated, image analysis was performed using ImageJ v1.54f. Initially, the Z-stacks were converted into 8-bit and then to a maximum intensity Z-projection image. Macros were used for quantitative analysis to ensure standardization across all samples whenever possible. For quantification of the fluorescent area, brightness/contrast adjustments were performed, if necessary, before thresholding the image and measuring the area. For quantification of objects (i.e., number of nuclei), following image thresholding, the “Analyze Particles” function was applied, along with the “Watershed” and “Open” corrections to separate merged objects. For dextran penetration, the area of the organoid tissue was manually outlined based on the nuclear staining.

### Morphological analysis of endothelial networks

3D projections of the whole organoids were obtained using ImageJ, as mentioned above. To quantify the percentage of CD31+ area, the total area of the organoid was calculated by measuring the largest βIIITUB+ object under saturation conditions. Then, CD31+ fluorescent area was normalized to its respective organoid total area. For the morphological analysis of the endothelial networks, binary masks were generated from the CD31+ channel. The masks were processed using the “Mexican hat filter” plugin, and from these images, the diameter of the networks was calculated using the command “Geometry to Distance Map” and the plugin “Diameter Measurements”. The same processed images were analyzed using AngioTool64 v0.6a for a more detailed morphological analysis of the networks.

### Statistical analysis and data visualization

Each batch of organoids was considered an independent repeat, and each individual organoid was considered a replicate or N. Two different iPSCs lines (A– and S-iPSCs) were tested in all experiments to increase the statistical power and highlight the differences between two cell lines that behave remarkably differently in terms of endogenous endothelial differentiation. If not explicitly stated, for most experiments, four or three batches of each iPSC line were used, with an average of four organoids per condition and per batch analyzed. All graphical representations and statistical analyses were performed using GraphPad Prism v8.0.2. The type of data visualization graph was chosen depending on the emphasized aspect (i.e., scatter plots to show individual variability or box/violins to show distribution). Data are presented as mean ± standard deviation (SD) unless for violin plots, in which data is presented as data distribution (violin shape), median and quartiles. For comparison between two groups, an unpaired t-test was used for normally distributed data with equal variances, Welch’s t-test when variances were unequal, and a Mann-Whitney test for non-normally distributed data. For multiple group comparisons, a one-way ANOVA followed by Tukey’s post hoc comparisons was used for normally distributed data with equal variances, Brown-Forsythe and Welch ANOVA with Games-Howell’s post hoc comparisons when variances were unequal, and a Kruskal-Wallis test followed by Dunn’s post hoc test for non-normally distributed data. For comparison with multiple variables, a two-way ANOVA was used with Tukey’s or Sidak’s post hoc comparisons. Details on the number of organoids (N), number of organoid batches, cell lines used, statistical tests and p-value legend can be found in the footnotes of each figure.

## ACKNOWLEDGMENTS

We thank Dr. Meenakshi Suku (Trinity College Dublin), Prof. Michael Monaghan (Trinity College Dublin) and Prof. Lesley Forester (University of Edinburgh) for facilitating the use of Zombie Green iPSCs, Dr. Mary Ní Fhlathartaigh and Prof. John Dalton for the use of the EVOS M5000 imaging system and Dr. Olena Kudina for her guidance during the nanoindentation measurements. We also express our gratitude to Prof. Dr. med. Sven Hendrix (MSH Medical School Hamburg – University of Applied Sciences and Medical University) for providing feedback on the manuscript.

The authors acknowledge the facilities and scientific and technical assistance of the Anatomy Imaging and Microscopy Facility (Prof. Peter Dockery, Dr. Peter Owens, https://imaging.universityofgalway.ie/imaging/), the Histology Suite (Dr. Keith Madden), the High throughput screening facility (Dr. Enda O’Connell, https://www.universityofgalway.ie/remedi/facilities/) and the Flow Cytometry Core Facility (Dr. Shirley Hanley, https://www.universityofgalway.ie/our-research/facilities/).

This work was supported by CÚRAM, Research Ireland Centre for Medical Devices, the School of Biological and Chemical Sciences, College of Science and Engineering, University of Galway, the Research Ireland 18/EPSRC-CDT/3583, Engineering and Physical Sciences Research Council EP/S02347X/1, European Union’s Horizon Europe Excellent Science programme under the Marie Skłodowska-Curie Actions Grant Agreement (Grant Agreement No 101081457), Research Ireland and the European Regional Development Fund (ERDF) under grant number 13/RC/2073_P2, and Simons Initiative for the Developing Brain (grant number 529085).

## AUTHOR CONTRIBUTIONS

**Josep Fumadó Navarro:** Conceptualization, Investigation, Data Curation, Formal analysis, Visualization, Methodology, Writing– original draft, Writing– review & editing. **Siobhan Crilly:** Methodology, Investigation, Writing– original draft, Writing– review & editing. **Wai Kit Chan, John O. Mason, Shane Browne, Catalina Vallejo Giraldo:** Methodology, Writing– review & editing. **Abhay Pandit:** Supervision, Resources, Methodology, Writing– review & editing. **Mihai Lomora**: Conceptualization, Methodology, Resources, Funding acquisition, Project administration, Supervision, Writing– original draft, Writing– review & editing.

## Supplemental Information

**Figure S1:**
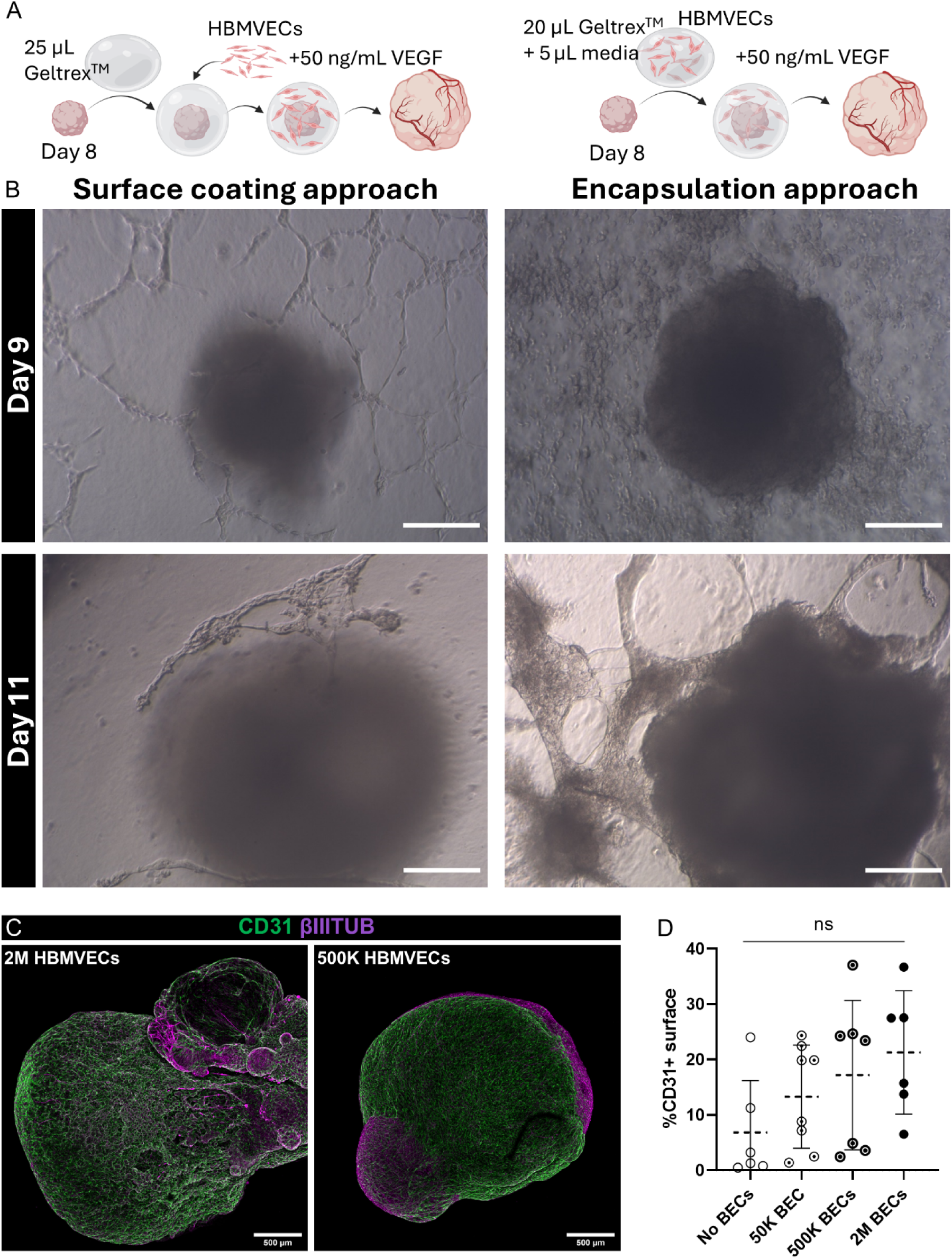
Optimization of the vascularization strategy by testing two different HBMVECs incorporation methods and varying cell densities. (A) Schematic representation of the surface coating approach consisting of adding HBMVECs after polymerizing the Geltrex^TM^ droplet embedding the organoid, allowing cell attachment (left panel). The encapsulation approach involved mixing HBMVECs with the biomaterial before embedding the organoid, followed by polymerization (right). (B) Brightfield images showing 50,000 HBMVECs-organoid interactions in the first days after embedding (days 9 and 11) for both approaches. In the surface coating approach, endothelial networks are formed on the surface but deteriorate over time. In contrast, the encapsulation approach resulted in more stable network assemblies. Scale bars: 500 µm. (C) Confocal images of A-COs grown with two different encapsulating cell densities (500,000 and 2 million HBMVECs), whole-stained for CD31 (green) and βIIITUB (magenta), depicting the organoid surface covered with HMBVECs. Scale bars: 500 µm. (D) Percentage of CD31+ coverage on day 40 of A-COs grown with different encapsulating cell densities (0, 50,000, 500,000 and 2 million HBMVECs). Mean ± SD, N = 6-8 organoids, two independent batches, same A-iPSC line. One-way ANOVA test and Tukey’s post hoc comparisons; ns = no significant difference (p > 0.05).

**Figure S2:**
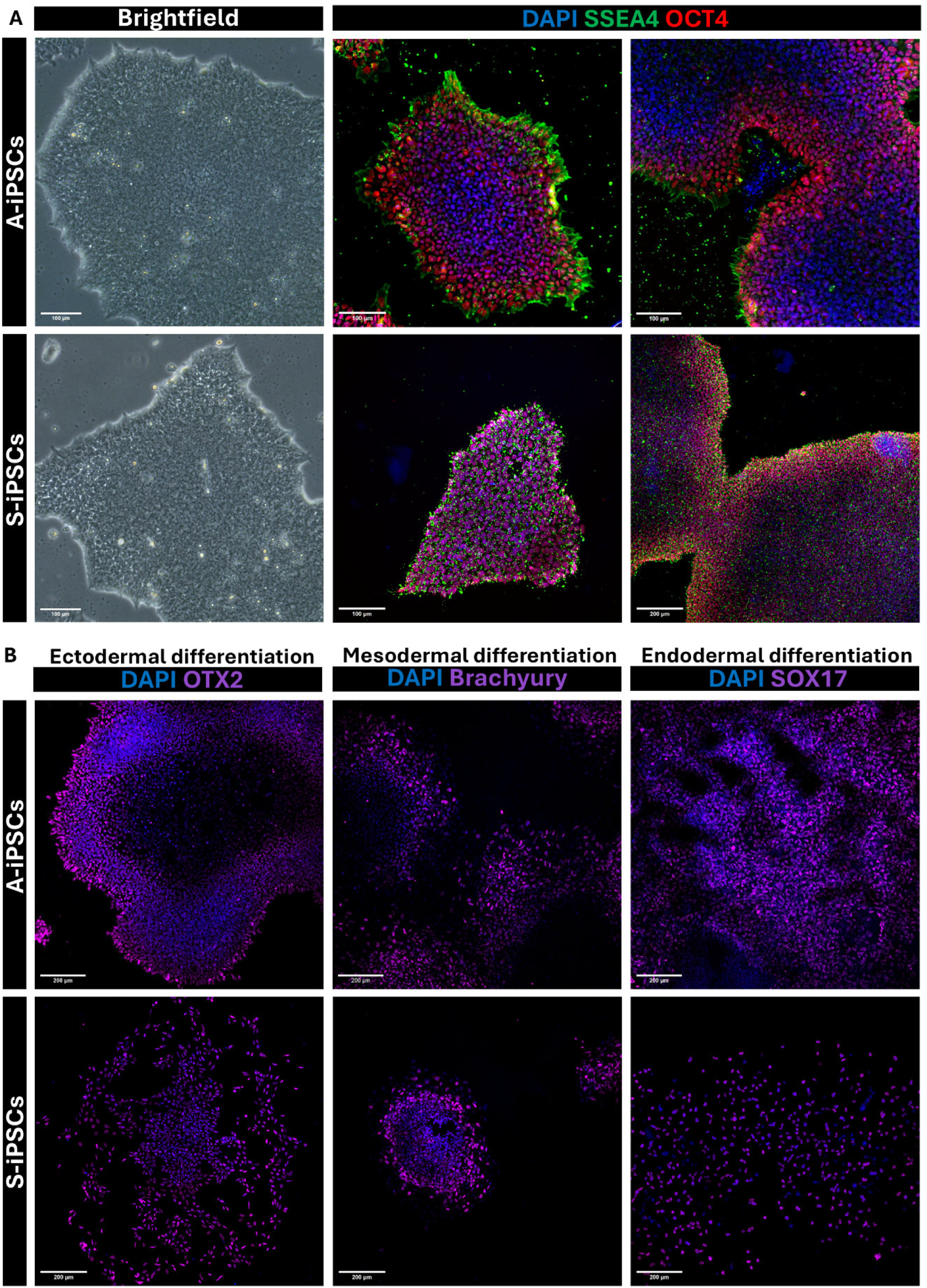
Quality control assessment of two iPSC lines (A and S) through colony morphology evaluation, staining for pluripotent markers and a three germinal layer differentiation assay. (A) Brightfield images of high-quality A– and S-iPSC cultures displaying compact colonies with well-defined edges. Confocal images of undifferentiated cultures stained for the pluripotency markers SSEA-4 (green), OCT4 (red), and nuclei (blue). Scale bars: 100 µm (bottom right image, 200 µm). (B) Representative confocal images of ectodermal, mesodermal and endodermal progenitors (left to right) derived from A– and S-iPSC lines after a three-germinal-layer differentiation assay. Cells were immunostained for lineage-specific markers: OTX2 (ectoderm), brachyury (mesoderm), and SOX17 (endoderm). Scale bars: 200 µm

**Figure S3:**
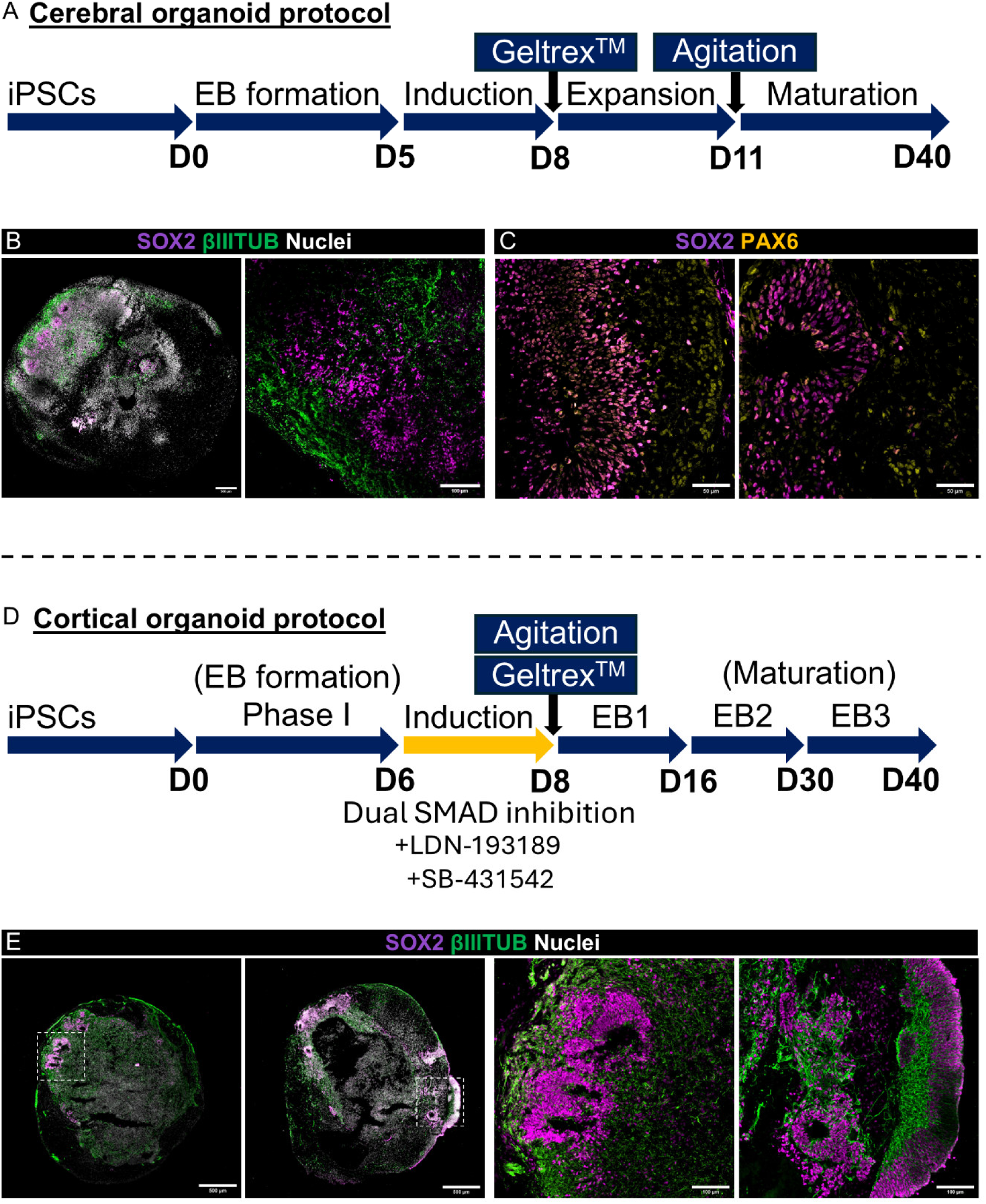
Comparison of the cerebral organoid (CO) and cortical organoid (CortiCO) protocols used in this study, and examples of typical brain organoid features obtained with both methods. (A) Schematic representation of key stages in the cerebral organoid protocol. (B) Confocal images of standard CO sections immunostained for SOX2 (neural stem cells, magenta), βIIITUB (neurons, green), and nuclei (white). Scale bars: 500 µm (left) and 100 µm (right). (C) Confocal images of standard CO sections, showing neural rosettes immunostained for SOX2 (neural stem cells, magenta), PAX6 (radial glia, yellow), and nuclei (white). Scale bars: 50 µm. (D) Schematic representation of key stages in the cortical organoid protocol. (E) Confocal images of standard CortiCOs sections immunostained for SOX2 (neural stem cells, magenta), βIIITUB (neurons, green), and nuclei (white). Scale bars: 500 µm (left) and 100 µm (right).

**Figure S4:**
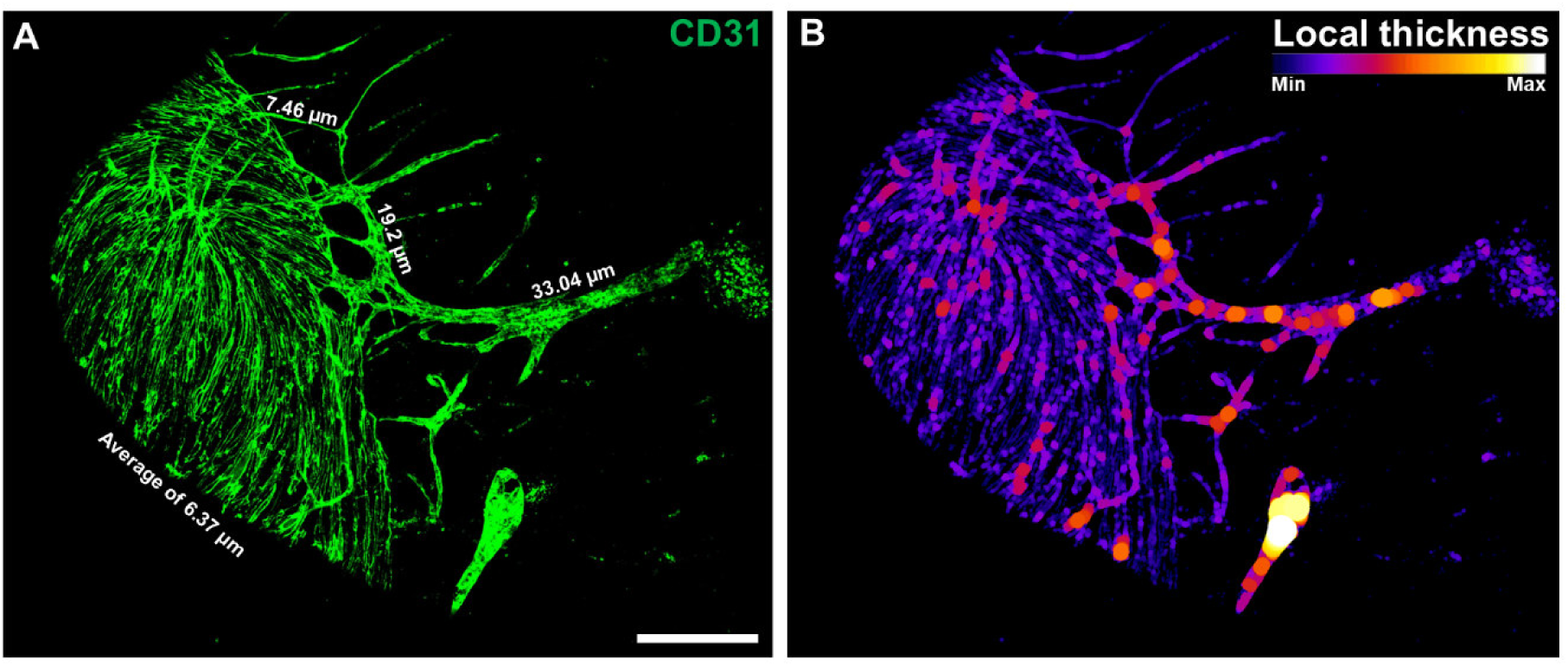
Evidence of hierarchical morphology in some endothelial networks. (A) Confocal image of a hierarchical endothelial network (CD31+) within a vascularized CO, showing a gradual decrease in vessel diameter from the right to the left. Scale bar: 200 µm. (B) Visualization of vessel diameter variations using the Local Thickness plugin of ImageJ.

**Figure S5:**
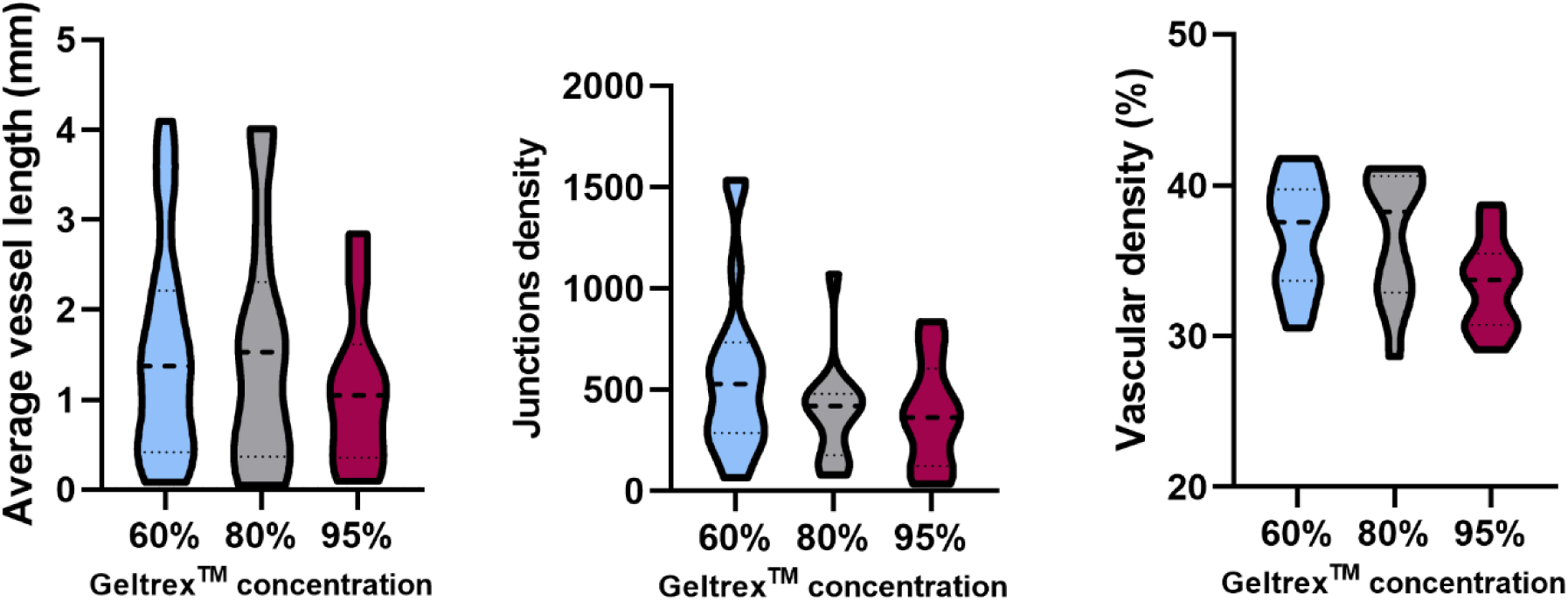
Additional morphological analysis of superficial endothelial networks formed within different Geltrex^TM^ concentrations. Quantification of junction density, average vessel length (mm), and vascular density (%) of superficial endothelial networks formed on organoids encapsulated with different Geltrex^TM^ concentrations. Light blue (60%), white (80%), Bordeaux (95%). Violin shape represents data distribution, Median (thick line), interquartile range (thin lines), N = 9-22, five different batches, same A-iPSC line. One-way ANOVA test and Tukey’s post hoc comparisons; not significant (p > 0.05).

**Figure S6:**
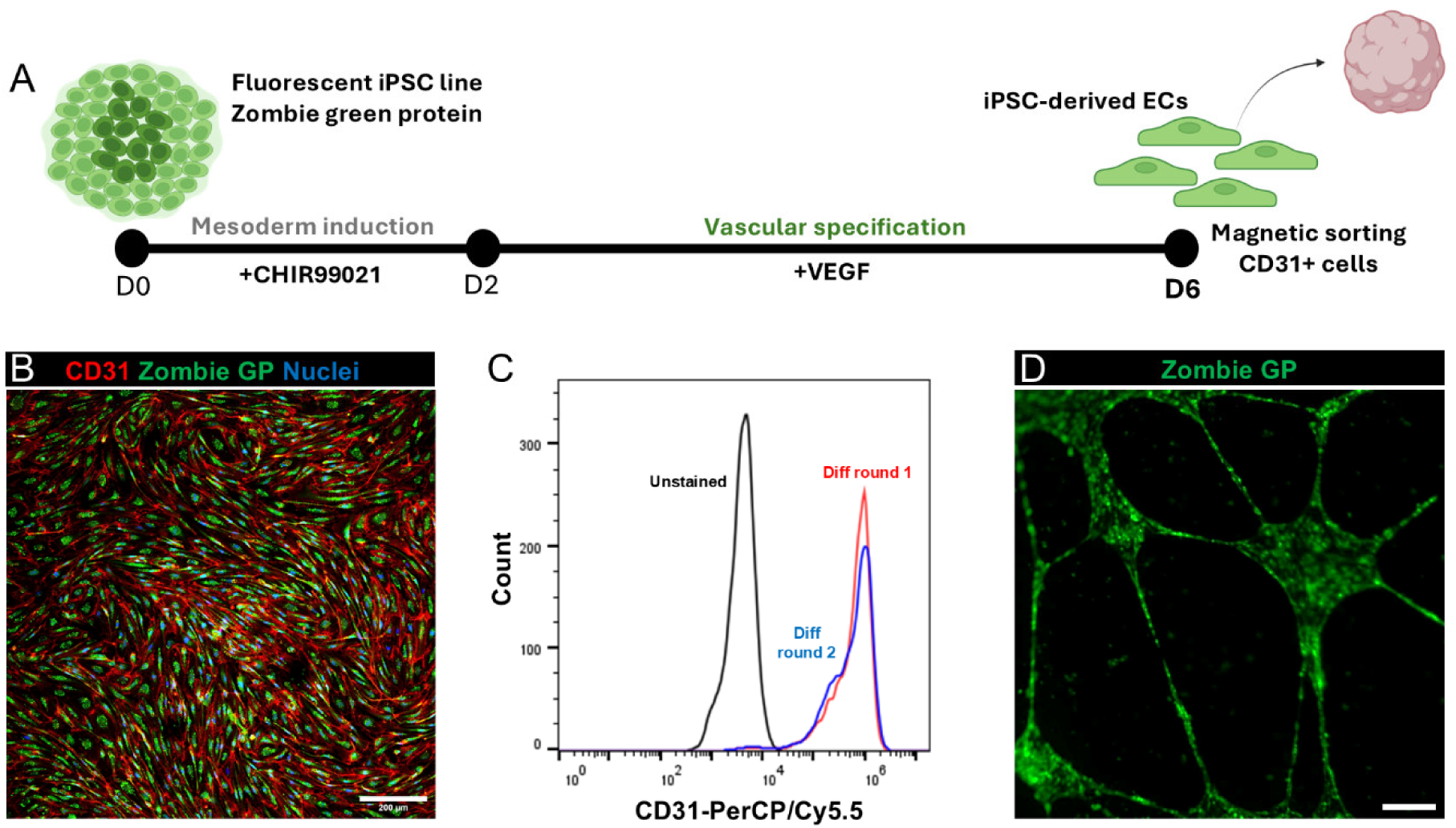
Generation and characterization of fluorescent iPSC-derived endothelial cells (Z-ECs) (A) Schematic representation of the iPSC-to-endothelial cell differentiation protocol. Created with Biorender.com. (B) Confocal image of fluorescent iPSC-derived endothelial cells (Z-ECs) expressing Zombie Green protein in the cytoplasm, post-sorting. Cells were immunostained for CD31 (red) and nuclei (blue). Scale bar: 200 µm. (C) Flow cytometry histogram of iPSC-derived ECs multiple passages after magnetic sorting. Black line, unstained cells; red and blue lines, two independent differentiations of cells labelled with a PerCP/Cy5.5-conjugated anti-CD31 antibody. (D) Fluorescence microscopy image of a Z-ECs tubular assay on 100% Geltrex^TM^. Scale bar: 200 µm.

**Figure S7:**
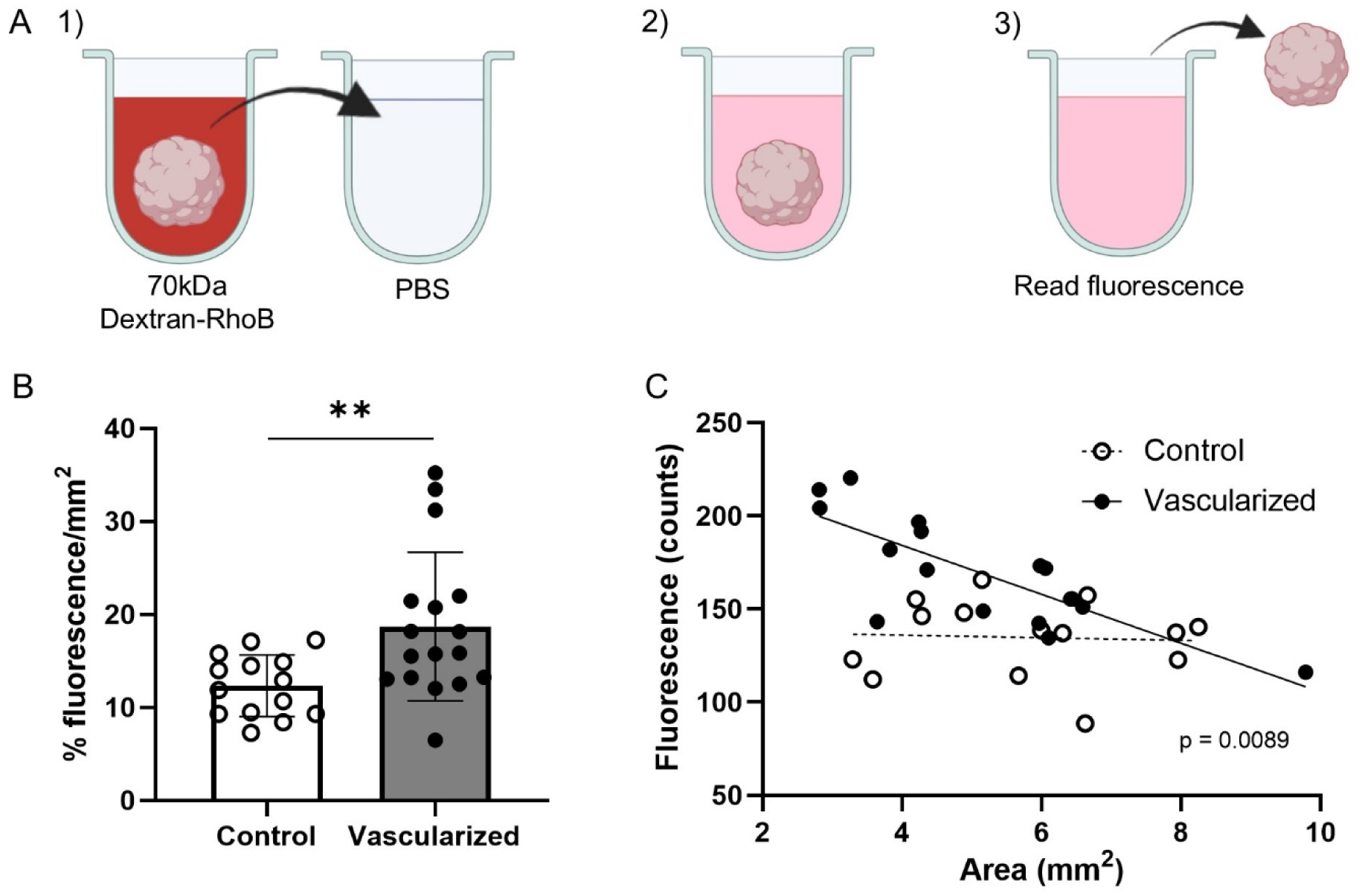
70 kDa dextran rhodamine B (d-RhoB) accumulation in control and vascularized COs and its correlation with organoid size. (A) Schematic representation of the experimental workflow. 1) Organoids were incubated in d-RhoB solution and transferred with the same volume of media to a well filled with PBS. 2) Organoids were incubated with shaking to release the fluorescent content. 3) Organoids were removed with the same volume prior to fluorescence measurements (;\_ex_ = 570,;\_ex_ = 595 nm). (B) Percentage of fluorescent counts transferred within the volume of d-RhoB containing the organoid, relative to the maximum fluorescence possible (same volume of d-RhoB solution alone), normalized by the area of the organoid (mm^2^). The same experiment was performed with control (white bar) and vascularized (grey bar) organoids from both A– and S-iPSC lines. Mean ± SD, N = 14-17, two independent batches, two iPSC lines. Welch’s T-test; **p < 0.01. (C) Individual organoid sizes (mm^2^) and their respective fluorescence values. White dots (control organoids), black dots (vascularized organoids), and black line shows the linear regression trend. N = 14-15 per conditions, two independent batches, two iPSC lines. Regressions are significantly different: p = 0.0089.

## REFERENCES

1. Durens M, Nestor J, Williams M, et al. High-throughput screening of human induced pluripotent stem cell-derived brain organoids. J Neurosci Methods. 2020;335:108627. doi:10.1016/J.JNEUMETH.2020.108627

2. Phan N, Hong JJ, Tofig B, et al. A simple high-throughput approach identifies actionable drug sensitivities in patient-derived tumor organoids. Commun Biol 2019 21. 2019;2(1):1–11. doi:10.1038/s42003-019-0305-x

3. Tang XY, Wu S, Wang D, et al. Human organoids in basic research and clinical applications. Signal Transduct Target Ther 2022 71. 2022;7(1):1–17. doi:10.1038/s41392-022-01024-9

4. Kim J, Koo BK, Knoblich JA. Human organoids: model systems for human biology and medicine. Nat Rev Mol Cell Biol 2020 2110. 2020;21(10):571–584. doi:10.1038/s41580-020-0259-3

5. Renner M, Lancaster MA, Bian S, et al. Self-organized developmental patterning and differentiation in cerebral organoids. EMBO J. 2017;36(10):1316–1329. doi:10.15252/embj.201694700

6. Arlotta P, Paşca SP. Cell diversity in the human cerebral cortex: from the embryo to brain organoids. Curr Opin Neurobiol. 2019;56:194–198. doi:10.1016/j.conb.2019.03.001

7. Mayhew CN, Singhania R. A review of protocols for brain organoids and applications for disease modeling. STAR Protoc. 2023;4(1):101860. doi:10.1016/J.XPRO.2022.101860

8. Eigenhuis KN, Somsen HB, van der Kroeg M, et al. A simplified protocol for the generation of cortical brain organoids. Front Cell Neurosci. 2023;17:1114420. doi:10.3389/FNCEL.2023.1114420/BIBTEX

9. Lancaster MA, Knoblich JA. Generation of cerebral organoids from human pluripotent stem cells. Nat Protoc. 2014;9(10):2329–2340. doi:10.1038/nprot.2014.158

10. Gabriel E, Gopalakrishnan J. Generation of iPSC-derived Human Brain Organoids to Model Early Neurodevelopmental Disorders. J Vis Exp. 2017;2017(122):55372. doi:10.3791/55372

11. Susaimanickam PJ, Kiral FR, Park I-H. Region Specific Brain Organoids to Study Neurodevelopmental Disorders. Int J stem cells. 2022;15(1):26–40. doi:10.15283/IJSC22006

12. Kadoshima T, Sakaguchi H, Nakano T, et al. Self-organization of axial polarity, inside-out layer pattern, and species-specific progenitor dynamics in human ES cell-derived neocortex. Proc Natl Acad Sci U S A. 2013;110(50):20284–20289. doi: 10.1073/pnas.1315710110

13. Pellegrini L, Bonfio C, Chadwick J, Begum F, Skehel M, Lancaster MA. Human CNS barrier-forming organoids with cerebrospinal fluid production. Science (80-). 2020;369(6500). doi:10.1126/science.aaz5626

14. Xiang Y, Tanaka Y, Cakir B, et al. hESC-Derived Thalamic Organoids Form Reciprocal Projections When Fused with Cortical Organoids. Cell Stem Cell. 2019;24(3):487–497.e7. doi:10.1016/J.STEM.2018.12.015

15. Muguruma K, Nishiyama A, Kawakami H, Hashimoto K, Sasai Y. Self-organization of polarized cerebellar tissue in 3D culture of human pluripotent stem cells. Cell Rep. 2015;10(4):537–550. doi:10.1016/J.CELREP.2014.12.051

16. Tieng V, Stoppini L, Villy S, Fathi M, Dubois-Dauphin M, Krause KH. Engineering of midbrain organoids containing long-lived dopaminergic neurons. Stem Cells Dev. 2014;23(13):1535–1547. doi:10.1089/SCD.2013.0442

17. Rosebrock D, Arora S, Mutukula N, et al. Enhanced cortical neural stem cell identity through short SMAD and WNT inhibition in human cerebral organoids facilitates emergence of outer radial glial cells. Nat Cell Biol 2022 246. 2022;24(6):981–995. doi:10.1038/s41556-022-00929-5

18. Zhang Z, O’Laughlin R, Song H, Ming G li. Patterning of brain organoids derived from human pluripotent stem cells. Curr Opin Neurobiol. 2022;74:102536. doi:10.1016/J.CONB.2022.102536

19. Li Y, Zeng PM, Wu J, Luo ZG. Advances and Applications of Brain Organoids. Neurosci Bull. 2023;39(11):1703. doi:10.1007/S12264-023-01065-2

20. Struzyna LA, Watt ML. The Emerging Role of Neuronal Organoid Models in Drug Discovery: Potential Applications and Hurdles to Implementation. Mol Pharmacol. 2021;99(4):256–265. doi:10.1124/MOLPHARM.120.000142

21. Kim SH, Chang MY. Application of Human Brain Organoids—Opportunities and Challenges in Modeling Human Brain Development and Neurodevelopmental Diseases. Int J Mol Sci 2023, Vol 24, Page 12528. 2023;24(15):12528. doi:10.3390/IJMS241512528

22. Wray S. Modelling neurodegenerative disease using brain organoids. Semin Cell Dev Biol. 2021;111:60–66. doi:10.1016/J.SEMCDB.2020.05.012

23. LaMontagne E, Muotri AR, Engler AJ. Recent advancements and future requirements in vascularization of cortical organoids. Front Bioeng Biotechnol. 2022;10:1048731. doi:10.3389/FBIOE.2022.1048731/BIBTEX

24. Qian X, Song H, Ming GL. Brain organoids: advances, applications and challenges. Development. 2019;146(8). doi:10.1242/DEV.166074

25. Bhaduri A, Andrews MG, Mancia Leon W, et al. Cell stress in cortical organoids impairs molecular subtype specification. Nat 2020 5787793. 2020;578(7793):142–148. doi:10.1038/s41586-020-1962-0

26. Vallon M, Chang J, Zhang H, Kuo CJ. Developmental and pathological angiogenesis in the central nervous system. Cell Mol Life Sci C. 2014;71(18):3489. doi:10.1007/S00018-014-1625-0

27. Konan LM, Reddy V, Mesfin FB. Neuroanatomy, Cerebral Blood Supply. StatPearls. Published online July 24, 2023. Accessed January 12, 2025. https://www.ncbi.nlm.nih.gov/books/NBK532297/

28. Clarke DD, Sokoloff Donald Dudley Clarke L, Sokoloff L, Dudley D., Circulation and energy metabolism in the brain. 1999;81. / Donald D. Clarke and Louis Sokoloff” (1999). Chemistry Faculty Publications 81. Accessed February 28, 2025.

29. Iadecola C. The Neurovascular Unit Coming of Age: A Journey through Neurovascular Coupling in Health and Disease. Neuron. 2017;96(1):17–42. doi:10.1016/J.NEURON.2017.07.030

30. Patabendige A, Janigro D. The role of the blood-brain barrier during neurological disease and infection. Biochem Soc Trans. 2023;51(2):613–626. doi:10.1042/BST20220830

31. Barar J, Rafi MA, Pourseif MM, Omidi Y. Blood-brain barrier transport machineries and targeted therapy of brain diseases. Bioimpacts. 2016;6(4):225–248. doi:10.15171/BI.2016.30

32. Daneman R, Prat A. The Blood – Brain Barrier. Cold Spring Harb Perspect Biol. 2015 Jan 5;7(1):a020412. doi: 10.1101/cshperspect.a02041

33. Mennen RH, Oldenburger MM, Piersma AH. Endoderm and mesoderm derivatives in embryonic stem cell differentiation and their use in developmental toxicity testing. Reprod Toxicol. 2022;107:44–59. doi:10.1016/J.REPROTOX.2021.11.009

34. Sahu S, Sharan SK. Translating Embryogenesis to Generate Organoids: Novel Approaches to Personalized Medicine. iScience. 2020;23(9):101485. doi:10.1016/J.ISCI.2020.101485

35. Cakir B, Xiang Y, Tanaka Y, et al. Engineering of human brain organoids with a functional vascular-like system. Nat Methods 2019 1611. 2019;16(11):1169–1175. doi:10.1038/s41592-019-0586-5

36. Ham O, Jin YB, Kim J, Lee MO. Blood vessel formation in cerebral organoids formed from human embryonic stem cells. Biochem Biophys Res Commun. 2020;521(1):84–90. doi:10.1016/j.bbrc.2019.10.079

37. Shi Y, Sun L, Wang M, et al. Vascularized human cortical organoids (vOrganoids) model cortical development in vivo. PLOS Biol. 2020;18(5):e3000705. doi:10.1371/JOURNAL.PBIO.3000705

38. Pham MT, Pollock KM, Rose MD, et al. Generation of human vascularized brain organoids. Neuroreport. 2018;29(7):588–593. doi:10.1097/WNR.0000000000001014

39. Chai YC, To SK, Simorgh S, et al. Spatially Self-Organized Three-Dimensional Neural Concentroid as a Novel Reductionist Humanized Model to Study Neurovascular Development. Adv Sci. 2024;11(5):2304421. doi:10.1002/ADVS.202304421

40. Nzou G, Wicks RT, Wicks EE, et al. Human Cortex Spheroid with a Functional Blood Brain Barrier for High-Throughput Neurotoxicity Screening and Disease Modeling. Sci Reports 2018 81. 2018;8(1):1–10. doi:10.1038/s41598-018-25603-5

41. Song L, Yuan X, Jones Z, et al. Assembly of Human Stem Cell-Derived Cortical Spheroids and Vascular Spheroids to Model 3-D Brain-like Tissues. Sci Rep. 2019;9(1). doi:10.1038/S41598-019-42439-9

42. Kook MG, Lee S-EE, Shin N, et al. Generation of Cortical Brain Organoid with Vascularization by Assembling with Vascular Spheroid. Int J stem cells. 2022;15(1):85–94. doi:10.15283/IJSC21157

43. Crouch EE, Bhaduri A, Andrews MG, et al. Ensembles of endothelial and mural cells promote angiogenesis in prenatal human brain. Cell. 2022;185(20):3753–3769.e18. doi:10.1016/j.cell.2022.09.004

44. Sun XY, Ju XC, Li Y, et al. Generation of Vascularized Brain Organoids to Study Neurovascular Interactions. Elife. 2022;11:1–28. doi:10.7554/eLife.76707

45. Ahn Y, An JH, Yang HJ, et al. Human Blood Vessel Organoids Penetrate Human Cerebral Organoids and Form a Vessel-Like System. Cells. 2021;10(8). doi:10.3390/CELLS10082036

46. Salmon I, Grebenyuk S, Abdel Fattah AR, et al. Engineering neurovascular organoids with 3D printed microfluidic chips. Lab Chip. 2022. doi:10.1039/D1LC00535A

47. Zhu Y, Sun L, Fu X, et al. Engineering microcapsules to construct vascularized human brain organoids. Chem Eng J. 2021;424:130427. doi:10.1016/J.CEJ.2021.130427

48. Cadena MA, Sing A, Taylor K, et al. A 3D Bioprinted Cortical Organoid Platform for Modeling Human Brain Development. Adv Healthc Mater. 2024;13(27):2401603. doi:10.1002/ADHM.202401603

49. Browne S, Gill EL, Schultheiss P, Goswami I, Healy KE. Stem cell-based vascularization of microphysiological systems. Stem Cell Reports. 2021;16(9):2058–2075. doi:10.1016/J.STEMCR.2021.03.015

50. Bernas MJ, Cardoso FL, Daley SK, et al. Establishment of primary cultures of human brain microvascular endothelial cells to provide an in vitro cellular model of the blood-brain barrier. Nat Protoc. 2010;5(7):1265–1272. doi:10.1038/NPROT.2010.76

51. Heath DE, Cooper SL. The development of polymeric biomaterials inspired by the extracellular matrix. J Biomater Sci Polym Ed. 2017;28(10-12):1051–1069. doi:10.1080/09205063.2017.1297285

52. Pepper MS. Role of the matrix metalloproteinase and plasminogen activator-plasmin systems in angiogenesis. Arterioscler Thromb Vasc Biol. 2001;21(7):1104–1117. doi:10.1161/HQ0701.093685

53. Das A, Fanslow W, Cerretti D, Warren E, Talarico N, McGuire P. Angiopoietin/Tek interactions regulate mmp-9 expression and retinal neovascularization. Lab Invest. 2003;83(11):1637–1645. doi:10.1097/01.LAB.0000097189.79233.D8

54. Wang H, Keiser JA. Vascular endothelial growth factor upregulates the expression of matrix metalloproteinases in vascular smooth muscle cells: role of flt-1. Circ Res. 1998;83(8):832–840. doi:10.1161/01.RES.83.8.832

55. Su Z, Zhang Y, Liao B, et al. Antagonism between the transcription factors NANOG and OTX2 specifies rostral or caudal cell fate during neural patterning transition. J Biol Chem. 2018;293(12):4445. doi:10.1074/JBC.M117.815449

56. Chen C, Rengarajan V, Kjar A, Huang Y. A matrigel-free method to generate matured human cerebral organoids using 3D-Printed microwell arrays. Bioact Mater. 2020;6(4):1130. doi:10.1016/J.BIOACTMAT.2020.10.003

57. Pagliaro A, Artegiani B, Hendriks D. Emerging approaches to enhance human brain organoid physiology. Trends Cell Biol. 2025;0(0):17. doi:10.1016/J.TCB.2024.12.001

58. Kozlowski MT, Crook CJ, Ku HT. Towards organoid culture without Matrigel. Commun Biol. 2021;4(1). doi:10.1038/S42003-021-02910-8

59. Martins-Costa C, Pham VA, Sidhaye J, et al. Morphogenesis and development of human telencephalic organoids in the absence and presence of exogenous extracellular matrix. EMBO J. 2023;42(22). doi:10.15252/EMBJ.2022113213

60. Wälchli T, Bisschop J, Carmeliet P, et al. Shaping the brain vasculature in development and disease in the single-cell era. Nat Rev Neurosci 2023 245. 2023;24(5):271–298. doi:10.1038/s41583-023-00684-y

61. Buonfiglioli A, Kübler R, Missall R, et al. A microglia-containing cerebral organoid model to study early life immune challenges. Brain Behav Immun. 2025;123:1127–1146. doi:10.1016/J.BBI.2024.11.008

62. Litvinov RI, Weisel JW. Fibrin mechanical properties and their structural origins. Matrix Biol. 2016;60-61:110. doi:10.1016/J.MATBIO.2016.08.003

63. Bupphathong S, Lim J, Fang HW, et al. Enhanced Vascular-like Network Formation of Encapsulated HUVECs and ADSCs Coculture in Growth Factors Conjugated GelMA Hydrogels. ACS Biomater Sci Eng. 2024;10(5):3306–3315. doi: 10.1021/acsbiomaterials.4c00465

64. Chen X, Liu C, McDaniel G, et al. Viscoelasticity of Hyaluronic Acid Hydrogels Regulates Human Pluripotent Stem Cell-derived Spinal Cord Organoid Patterning and Vascularization. Adv Healthc Mater. 2024;13(32):e2402199. doi:10.1002/ADHM.202402199

65. Aisenbrey EA, Murphy WL. Synthetic alternatives to Matrigel. Nat Rev Mater. 2020;5(7):539. doi:10.1038/S41578-020-0199-8

66. Stankovic I, Notaras M, Wolujewicz P, et al. Schizophrenia endothelial cells exhibit higher permeability and altered angiogenesis patterns in patient-derived organoids. Transl Psychiatry 2024 141. 2024;14(1):1–15. doi:10.1038/s41398-024-02740-2

67. Logan S, Arzua T, Yan Y, et al. Dynamic Characterization of Structural, Molecular, and Electrophysiological Phenotypes of Human-Induced Pluripotent Stem Cell-Derived Cerebral Organoids, and Comparison with Fetal and Adult Gene Profiles. Cells. 2020;9(5). doi:10.3390/CELLS9051301

68. Bruveris FF, Ng ES, Stanley EG, Elefanty AG. VEGF, FGF2, and BMP4 regulate transitions of mesoderm to endothelium and blood cells in a human model of yolk sac hematopoiesis. Exp Hematol. 2021;103:30–39.e2. doi:10.1016/J.EXPHEM.2021.08.006

69. Ding VMY, Ling L, Natarajan S, Yap MGS, Cool SM, Choo ABH. FGF-2 modulates Wnt signaling in undifferentiated hESC and iPS cells through activated PI3-K/GSK3β signaling. J Cell Physiol. 2010;225(2):417–428. doi:10.1002/JCP.22214

70. Bonney S, Harrison-Uy S, Mishra S, et al. Diverse Functions of Retinoic Acid in Brain Vascular Development. J Neurosci. 2016;36(29):7786–7801. doi:10.1523/JNEUROSCI.3952-15.2016

71. Pawlikowski B, Wragge J, Siegenthaler JA. Retinoic acid signaling in vascular development. Genesis. 2019;57(7-8):e23287. doi:10.1002/DVG.23287

72. Shibuya M. Vascular Endothelial Growth Factor (VEGF) and Its Receptor (VEGFR) Signaling in Angiogenesis: A Crucial Target for Anti– and Pro-Angiogenic Therapies. Genes Cancer. 2011;2(12):1097. doi:10.1177/1947601911423031

73. Wrighton KH. The different flavours of iPS cells. Nat Rev Genet 2017 187. 2017;18(7):394–394. doi:10.1038/nrg.2017.42

74. Volpato V, Webber C. Addressing variability in iPSC-derived models of human disease: guidelines to promote reproducibility. Dis Model Mech. 2020;13(1):dmm042317. doi:10.1242/DMM.042317

75. Fedele G, Cazzaniga A, Castiglioni S, et al. The presence of BBB hastens neuronal differentiation of cerebral organoids – The potential role of endothelial derived BDNF. Biochem Biophys Res Commun. 2022;626:30–37. doi:10.1016/J.BBRC.2022.07.112

76. Sato Y, Asahi T, Kataoka K. Integrative single-cell RNA-seq analysis of vascularized cerebral organoids. BMC Biol. 2023;21(1):1–17. doi:10.1186/s12915-023-01711-1

77. Rosenstein JM, Krum JM, Ruhrberg C. VEGF in the nervous system. Organogenesis. 2010;6(2):107–114. doi:10.4161/ORG.6.2.11687

78. Lopez-Yrigoyen M, Fidanza A, Cassetta L, et al. A human iPSC line capable of differentiating into functional macrophages expressing ZsGreen: a tool for the study and in vivo tracking of therapeutic cells. Philos Trans R Soc B Biol Sci. 2018;373(1750). doi:10.1098/RSTB.2017.0219

79. Zudaire E, Gambardella L, Kurcz C, Vermeren S. A computational tool for quantitative analysis of vascular networks. PLoS One. 2011;6(11). doi:10.1371/JOURNAL.PONE.0027385

80. Rosebrock D, Arora S, Mutukula N, et al. Enhanced cortical neural stem cell identity through short SMAD and WNT inhibition in human cerebral organoids facilitates emergence of outer radial glial cells. Nat Cell Biol 2022 246. 2022;24(6):981–995. doi:10.1038/s41556-022-00929-5

81. Browne S, Hossainy S, Healy K. Hyaluronic Acid Macromer Molecular Weight Dictates the Biophysical Properties and in Vitro Cellular Response to Semisynthetic Hydrogels. ACS Biomater Sci Eng. 2020;6(2):1135–1143. doi: 10.1021/acsbiomaterials.9b01419

82. Qi Y, Yu T, Xu J, et al. FDISCO: Advanced solvent-based clearing method for imaging whole organs. Sci Adv. 2019;5(1):eaau8355. doi:10.1126/SCIADV.AAU8355

